# On-demand, reversible blood-brain barrier opening via electrical activation of piezoelectric nanoparticles for targeted brain drug delivery

**DOI:** 10.64898/2026.07.20.738792

**Authors:** Salona Roy, Umisha Siwakoti, Daniel Alday, Carlos Astete, Ethan McElveen, Ashok Sigdel, Fabio Del Piero, Cristina Sabliov, Elisa Castagnola, Qi Cai

## Abstract

Delivering therapeutics to the brain remains one of the most persistent challenges in medicine, because the blood-brain barrier (BBB) excludes over 98% of small-molecule drugs and virtually all biologics from the central nervous system (CNS). We developed electrical BBB modulation (eBBB), an on-demand platform combining vascular-targeting poly-L-lactic acid nanoparticles with high-definition transcranial direct current stimulation to achieve spatially and temporally controlled BBB opening. eBBB produced localized, reversible increases in BBB permeability confined to the stimulated cortex, with the opening area tunable via electrode geometry. This transient window enhanced regional delivery of a small-molecule drug, full-length immunoglobulins, and adeno-associated viral vectors, which are cargo classes otherwise completely excluded by the intact BBB. Neurovascular unit architecture was preserved with no lasting histological damage. Integrating a biodegradable nanomaterial with a clinically evaluated stimulation technology, eBBB offers a programmable, minimally invasive strategy for regional CNS drug delivery across brain malignancies and neurological disorders.

**One sentence summary:** Electrical activation of piezoelectric nanoparticles reversibly opens the blood-brain barrier for minimally invasive drug delivery to targeted cortical regions.

## Introduction

The blood-brain barrier (BBB) is a highly selective and tightly regulated interface that protects the central nervous system (CNS) from blood-borne toxins and pathogens (*1*). Tight junction (TJ) complexes between adjacent brain endothelial cells (ECs) establish the physical basis of the barrier, while pericytes and astrocytic end-feet dynamically regulate its function; together with microglia and neurons, these elements form the neurovascular unit (NVU), a multicellular system whose coordinated signaling dynamically regulates barrier integrity in response to both physiological and pathological cues (*2–4*). Unlike peripheral ECs, brain ECs exhibit minimal transcytosis, limiting vesicle-mediated transport (*5–9*). A network of active efflux transporters further expels diverse substrates, including many therapeutic agents, back into the bloodstream against their concentration gradient (*10–13*). As a result, the BBB excludes over 98% of small-molecule drugs and virtually all biologics from the CNS, posing a fundamental obstacle to treating brain tumors, neurodegenerative diseases, and other neurological disorders (*14, 15*).

Current strategies to modulate the BBB either exploit endogenous transport pathways or transiently increase barrier permeability. Intracarotid injection of hyperosmotic mannitol induces EC shrinkage and TJ opening, enabling passive diffusion across the barrier, but produces broad BBB disruption with limited regional selectivity (*16, 17*). Receptor-mediated transcytosis has been explored using antibodies or peptides that engage endothelial transport systems, including the transferrin and insulin receptors. However, these approaches generally enhance delivery throughout the brain rather than within a defined anatomical region and must be adapted to the properties of each therapeutic cargo (*18–20*). Focused ultrasound combined with circulating microbubbles uses localized mechanical effects to transiently increase BBB permeability and has emerged as a leading strategy for regional brain delivery (*21, 22*). More recently, picosecond laser stimulation of vascular-targeted gold nanoparticles has been shown to mechanically modulate the BBB and enhance drug delivery in brain tumor models (*14, 23–25*). Nevertheless, there remains a need for a compact, programmable approach that can open the BBB in a selected brain region and adjust the spatial extent of the opening to the area requiring treatment. Such control may be particularly valuable for superficial brain metastases and frontocortical regions affected by neurodegenerative disease, which are located at or near the cortical surface (*26–28*).

Here, we introduce electrical BBB modulation (eBBB), a transcranial strategy that, for the first time, combines high-definition transcranial direct current stimulation (HD-tDCS) with the activation of albumin-functionalized poly-L-lactic acid (PLLA) nanoparticles. PLLA was chosen because it is biodegradable and has reported piezoelectric properties (*29*). Bovine serum albumin (Alb) functionalization was used to promote nanoparticle association with brain microvascular, leveraging its strong structural homology with human serum albumin and its ability to reduce non-specific protein adsorption in circulation (*30–32*). To achieve focal stimulation, custom multielectrode arrays (MEAs) were designed to concentrate the electrical field over a defined cortical region. We hypothesized that HD-tDCS would electromechanically activate endothelial-associated PLLA nanoparticles, generating localized mechanical stimuli at the vascular interface that transiently increase BBB permeability. By tuning nanoparticle dose, stimulation parameters, and electrode configuration, eBBB was designed to provide precise spatial and temporal control over BBB modulation.

We show that eBBB transiently reduces endothelial barrier resistance and increases permeability *in vitro*, whereas nanoparticles or electrical stimulation alone produce little effect. In mice, eBBB generates regionally confined and reversible BBB opening that can be tuned through nanoparticle dose, stimulation current, and stimulation duration. Moreover, the spatial extent of opening is adjustable through MEA montages. We further demonstrate that the eBBB platform successfully delivers the small molecule drug paclitaxel, immunoglobulins, and adeno-associated viral (AAV) vectors, cargo classes that normally cannot traverse the BBB, into the mouse brain parenchyma, establishing functional efficacy for both pharmacological and genetic intervention. NVU architecture was preserved, and no lasting histological damage was detected. Together, these results establish proof of concept for a programmable approach to delivering therapeutically distinct cargos across the BBB.

## Results

### PLLA-Alb nanoparticles exhibit enhanced endothelial association without altering baseline barrier function

To promote nanoparticle association with the brain microvascular endothelium, we prepared bovine serum albumin-conjugated PLLA nanoparticles (PLLA-Alb NPs) using a conjugate-first strategy, in which PLLA-Alb conjugates were formed prior to nanoparticle assembly. In parallel, PLLA-polyethylene glycol nanoparticles (PLLA-PEG NPs) were synthesized as a low- endothelial-association control. The PEG molecules (amine-PEG-carboxymethyl, 5 kDa) shield the surface from protein adsorption and are expected to reduce specific endothelial binding relative to albumin-coated particles (**Fig. 1A**) (*33*).

**Fig. 1.**
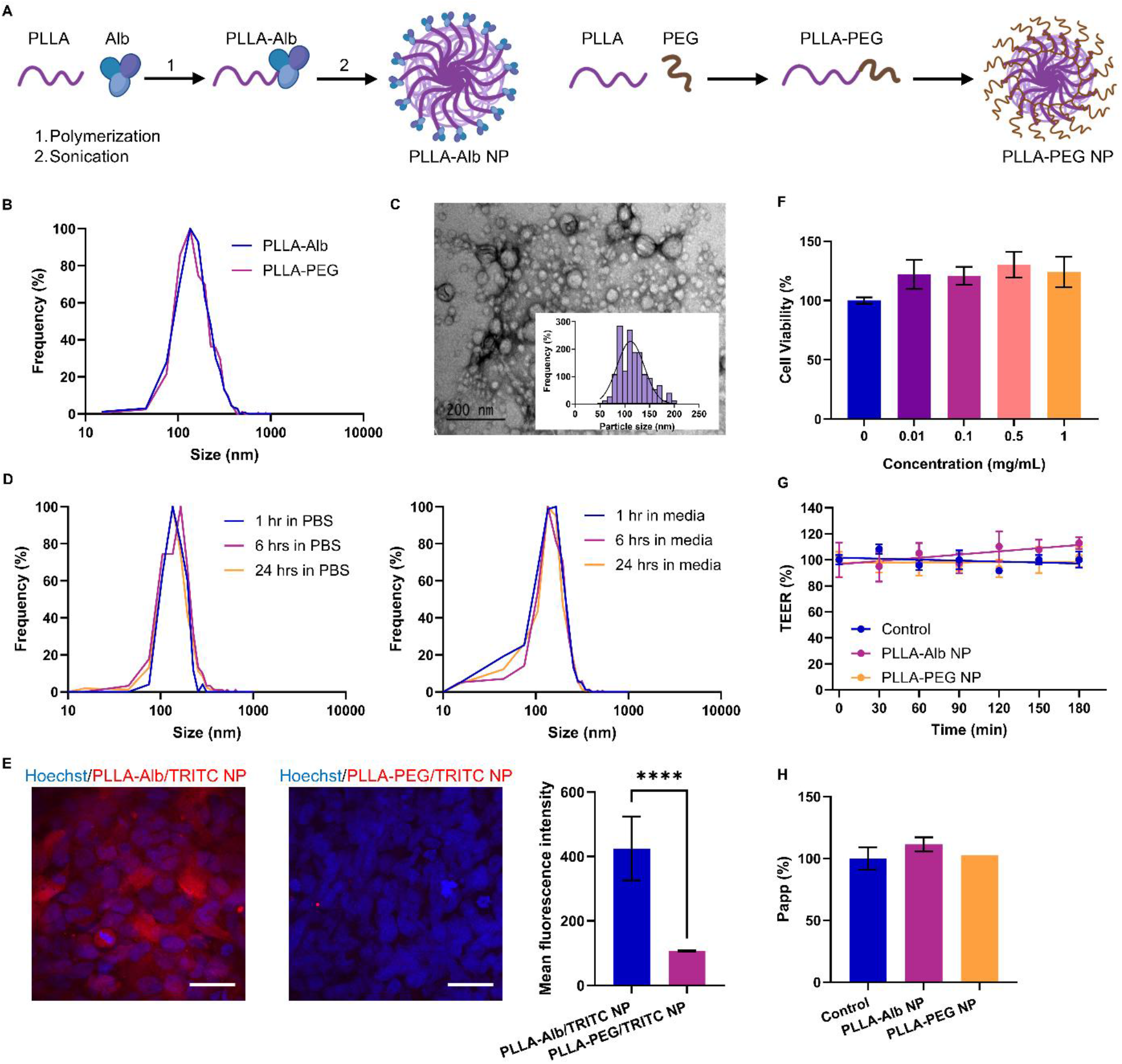
Synthesis and characterization of PLLA-Alb NPs for eBBB. (A) Synthesis workflow of PLLA-Alb and PLLA-PEG NPs. (B) DLS analysis showing the hydrodynamic diameters of PLLA-Alb and PLLA-PEG NPs. (C) TEM image of PLLA-Alb NP. The size distribution is shown in the inset histogram. 121 nanoparticles were measured using Image-J. (D) Colloidal stability assessment of PLLA-Alb NPs in PBS and cell culture media over 24 hours. (E) Fluorescence microscopy images showing selective uptake of PLLA-Alb/TRITC by hCMEC/D3 cells. Scale bar = 50 µm. (F) Cell viability of hCMEC/D3 cells treated with increasing concentrations (0.01–1 mg/mL) of PLLA-Alb NPs for 72 hours. (G) Assessment of nanoparticle biocompatibility in the *in vitro* BBB model by measuring the TEER. (H) Assessment of nanoparticle biocompatibility in the *in vitro* BBB model by measuring the apparent permeability to FITC–Dextran (40 kDa). For panels E–H, data are presented as mean ± SD (n = 3–6 biological replicates). Statistical analysis: (E) Unpaired Student’s two-sided *t*-test. ****P<0.0001. (F)–(H): One-way ANOVA, no significant differences.

The two formulations had nearly identical hydrodynamic diameters but distinct surface charges. PLLA-Alb and PLLA-PEG NPs had mean hydrodynamic diameters of 161 ± 70 nm and 161 ± 49 nm, respectively (**Fig. 1B**), and zeta potentials of −37.4 ± 0.5 mV and −15.6 ± 1.5 mV. Protein quantification indicated an albumin content of approximately 0.50 mg per mg of PLLA-Alb NPs.

TEM revealed that PLLA-Alb NPs were predominantly spherical, with an average diameter of 112 ± 29 nm (**Fig. 1C**); the smaller TEM diameter relative to the DLS hydrodynamic diameter is consistent with the collapsed dry state measured by electron microscopy compared with the hydrated particle in solution. Their hydrodynamic size changed little over 24 hours of incubation in PBS or cell culture medium supplemented with 5% fetal bovine serum, indicating colloidal stability under the experimental conditions **(Fig. 1D)**.

To determine whether albumin functionalization enhanced interaction with brain endothelial cells, we incubated human brain microvascular endothelial cell (hCMEC/D3) monolayers with TRITC-labeled PLLA-Alb or PLLA-PEG NPs for 30 minutes. PLLA-Alb NPs produced markedly greater cell-associated fluorescence than PLLA-PEG NPs (**Fig. 1E**). Confocal z-stack imaging further localized PLLA-Alb NPs to both the cell membrane and cytoplasm (**fig. S1**). Membrane-associated signal likely reflects surface binding, whereas cytoplasmic localization suggests partial internalization within the 30-minute incubation period. Having established enhanced endothelial association, we next examined whether the nanoparticles affected cell viability or baseline barrier function. PLLA-Alb NPs did not reduce WST-1 metabolic activity after 72 hours at concentrations ranging from 0.01 to 1 mg/mL (**Fig. 1F**). We then evaluated both nanoparticle formulations in an established human BBB Transwell model, in which hCMEC/D3 cells were cultured as a monoculture on the upper surface of the insert (**fig. S2A**) (*24*). Barrier properties were characterized by a transendothelial electrical resistance (TEER) of 14 ± 2 Ω·cm² and an apparent permeability to 40-kDa FITC-dextran of 9.16 ± 1.64 × 10⁻⁷ cm/s, with occludin expression confirmed by immunostaining and Western blot (**fig. S2, B to E**), consistent with published benchmarks for this cell line (*34, 35*). Although the TEER of this model is lower than that of primary brain endothelial cells, hCMEC/D3 monolayers have been widely adopted for BBB modulation studies owing to their human origin, stable barrier phenotype, and reproducibility. Following a 30-minute exposure to PLLA-Alb or PLLA-PEG NPs at 1 mg/mL, neither formulation significantly altered TEER or FITC-dextran permeability (**Fig. 1G** and H). Together, these results indicate that PLLA-Alb NPs were well tolerated and that neither nanoparticle formulation altered baseline barrier integrity, providing a stable foundation for subsequent studies of electrically induced BBB modulation.

### eBBB produces tunable and reversible BBB modulation *in vitro*

To characterize BBB permeability changes induced by eBBB, we designed custom tungsten (W) microelectrodes coated with nano-platinum (nano-Pt) to enable precise current delivery within the transwell insert. The wire microelectrodes were insulated with a thin layer of medical-grade epoxy, leaving a 4 mm segment at the tip intentionally exposed for electrical stimulation. The exposed tips were electrochemically coated with nano-Pt to increase the effective surface area (**Fig. 2, A and B**). Nano-Pt deposition was confirmed by scanning electron microscopy, which revealed a rough, nanostructured surface morphology consistent with increased charge injection capacity. The coated electrodes exhibited low impedance across the entire frequency range (10 Hz–100 kHz) (**Fig. 2C**), enabling safe, prolonged delivery of constant direct current while maintaining low and stable impedance (**Fig. 2D**) without inducing electrolysis, thereby preventing damage to either the electrodes or the cells. A pair of nano-Pt-coated electrodes (6 cm in length, 508 µm in diameter) was positioned on opposite sides of the transwell insert, with 4 mm of each coated tip submerged in the culture medium, thereby allowing direct electrical stimulation of the BBB model (**Fig. 3A**). We next varied the nanoparticle concentration (0.01–1 mg/mL), total stimulation current (0.02– 0.10 mA), and stimulation duration (1–5 minutes) to identify conditions that produced a reversible change in barrier function. At 0.5 mg/mL PLLA-Alb NPs, TEER decreased to approximately 60% of its baseline at 1 hour after stimulation and recovered by 3 hours (**Fig. 3B**). At 1 mg/mL, TEER did not fully recover within the 3-hour observation period, suggesting that excessive nanoparticle loading followed by electrical stimulation may prolong or irreversibly alter barrier remodeling, whereas lower concentrations produced little measurable change. Among the currents tested, 0.10 mA produced the most pronounced reversible response, and 5 minutes of stimulation was required to significantly reduce TEER (**Fig. 3, C and D**). Based on these results, we selected 0.5 mg/mL PLLA-Alb NPs, 0.10 mA, and 5 minutes of stimulation as the standard *in vitro* eBBB condition. Under these conditions, the apparent permeability of 40-kDa FITC-dextran increased 1.3-fold relative to the untreated control (**Fig. 3E**).

**Fig. 2.**
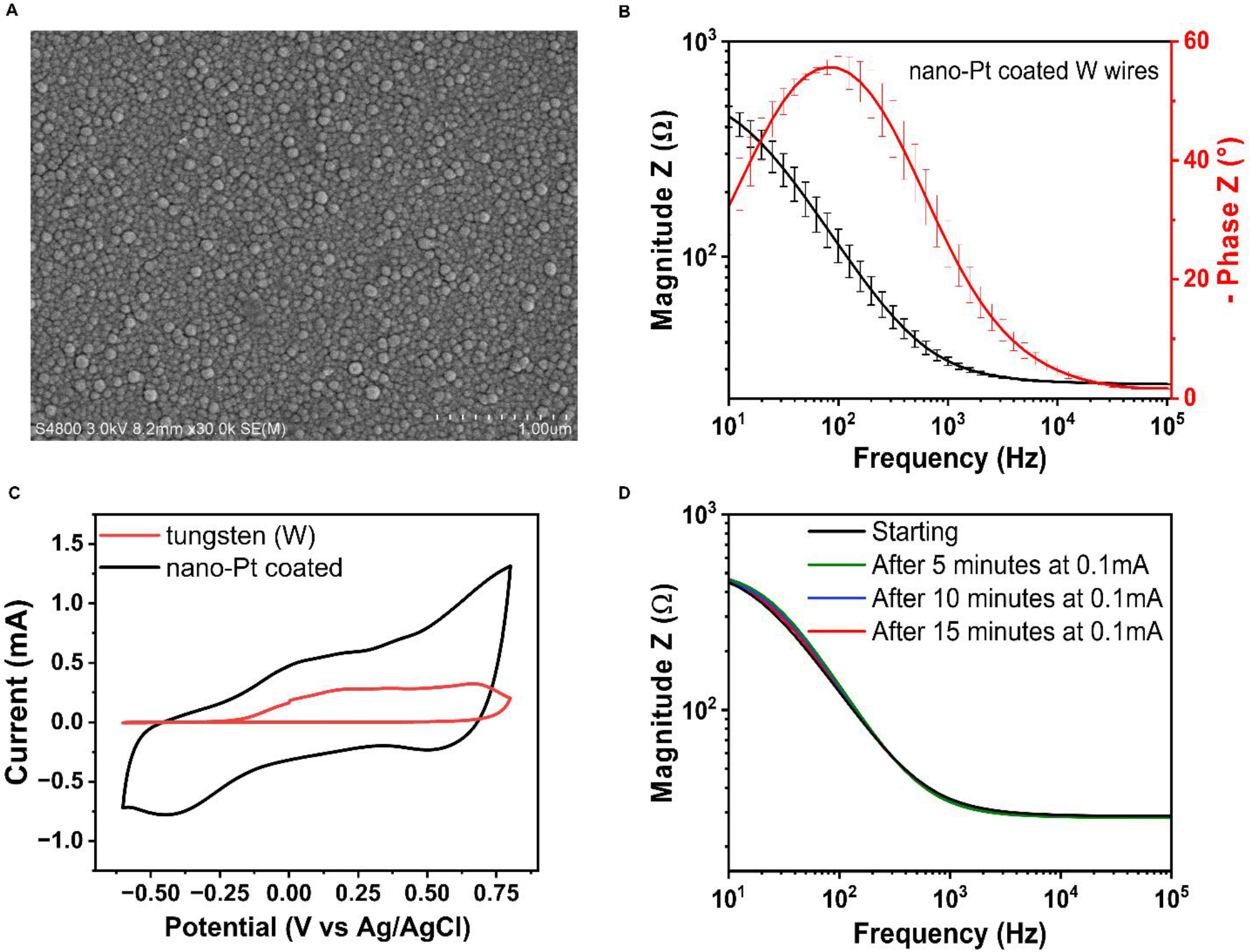
Morphological and electrochemical characterization of nano-Pt–coated tungsten wires for *in vitro* testing. (A) Scanning electron microscopy (SEM) image of the **n**ano-Pt coating showing the nanostructured surface morphology. (B) Electrochemical impedance spectroscopy (EIS; magnitude and phase angle) of nano-Pt-coated electrodes, showing low impedance across the high frequency range indicative of improved electrical conductivity and charge-transfer properties. (C) Cyclic voltammograms of bare tungsten and nano-Pt-coated wires, indicating increased surface area and enhanced charge transfer capability. (D) EIS spectra of nano-Pt-coated wires before and after application of a 0.1 mA constant current for 5, 10, and 15 minutes, demonstrating electrochemical stability under stimulation conditions.

**Fig. 3.**
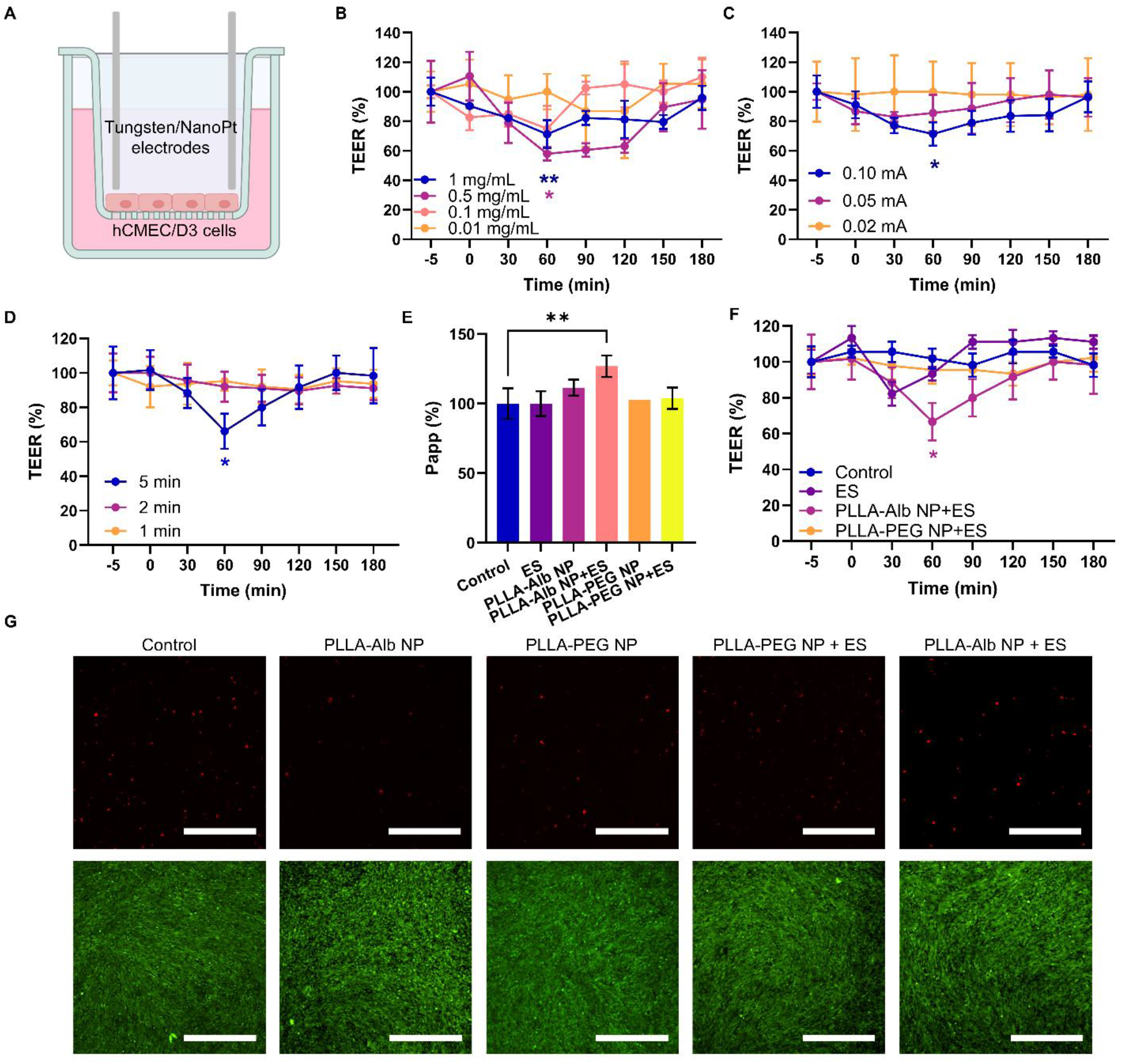
*In vitro* BBB modulation using eBBB. (A) Schematic illustration of the *in vitro* transwell BBB model equipped with custom tungsten/nano-platinum microelectrodes for focal electrical stimulation of PLLA-Alb NPs targeted at hCMEC/D3 cells. (B) Dose-dependent modulation of BBB permeability by PLLA-Alb NPs under fixed electrical stimulation (0.10 mA, 5 min). (C) Current-dependent modulation of BBB permeability by PLLA-Alb NPs under fixed nanoparticle dose (0.5 mg/mL) and electrical stimulation duration (5 minutes). (D) Time-dependent modulation of BBB permeability by PLLA-Alb NPs under fixed nanoparticle dose (0.5 mg/mL) and current (0.10 mA). (E) Changes in the apparent permeability of 40 kDa FITC-dextran across the BBB following eBBB treatment. ES: electrical stimulation only. (F) Changes in TEER following eBBB treatment. (G) Evaluation of eBBB safety using live/dead staining. Green indicates live cells, and red indicates dead cells. Scale bar = 100 µm. For panels (B), (C), (D), and (F), n = 3 biological replicates; for panel (E), n = 4 biological replicates. Statistical analysis: One-way ANOVA (*p < 0.05, **p < 0.01).

To determine whether the permeability response required both components of eBBB, we compared the combined treatment with electrical stimulation (ES) alone and with electrical stimulation following incubation with PLLA-PEG NPs. Neither control reproduced the decrease in TEER or increase in FITC-dextran permeability observed with PLLA-Alb NPs and electrical stimulation (**Fig. 3, E and F**). Together with the absence of barrier changes following nanoparticle exposure alone, these results show that eBBB depends on the combination of endothelial-associated PLLA- Alb NPs and electrical stimulation.

Live/dead fluorescence staining performed 1 hour after treatment revealed minimal cell death across all groups, including no-treatment control, PLLA-Alb NPs alone, PLLA-PEG NPs alone, PLLA-PEG NPs with ES, and eBBB (PLLA-Alb NPs with ES), confirming that neither the nanoparticles nor the stimulation parameters induced acute endothelial cytotoxicity (**Fig. 3G**). Thus, the optimized eBBB protocol produced a tunable and transient increase in endothelial barrier permeability without overt acute cell death.

### eBBB induces calcium-dependent paracellular barrier remodeling

To determine how eBBB increases endothelial permeability, we first examined calcium-dependent changes in endothelial junctions and the cytoskeleton. Intracellular Ca²⁺ is a well-established second messenger that regulates tight junction assembly and actomyosin contractility in brain endothelial cells; its elevation triggers myosin light chain kinase (MLCK) activation, leading to actin stress fiber formation and paracellular gap opening (*36, 37*). Pretreating hCMEC/D3 monolayers with BAPTA-AM, a cell-permeable intracellular Ca²⁺ chelator, progressively attenuated the eBBB-induced reduction in TEER in a dose-dependent manner. At 1 and 10 µM, BAPTA-AM largely prevented the response, whereas BAPTA-AM alone did not significantly alter baseline barrier resistance (**Fig. 4A** and **fig. S3**). BAPTA-AM was loaded during the 30-minute period following nanoparticle incubation and immediately before electrical stimulation, ensuring that Ca²⁺ was chelated at the time of electromechanical activation without interfering with nanoparticle association. These findings indicate that intracellular Ca²⁺ availability is required for eBBB-induced barrier modulation.

**Fig. 4.**
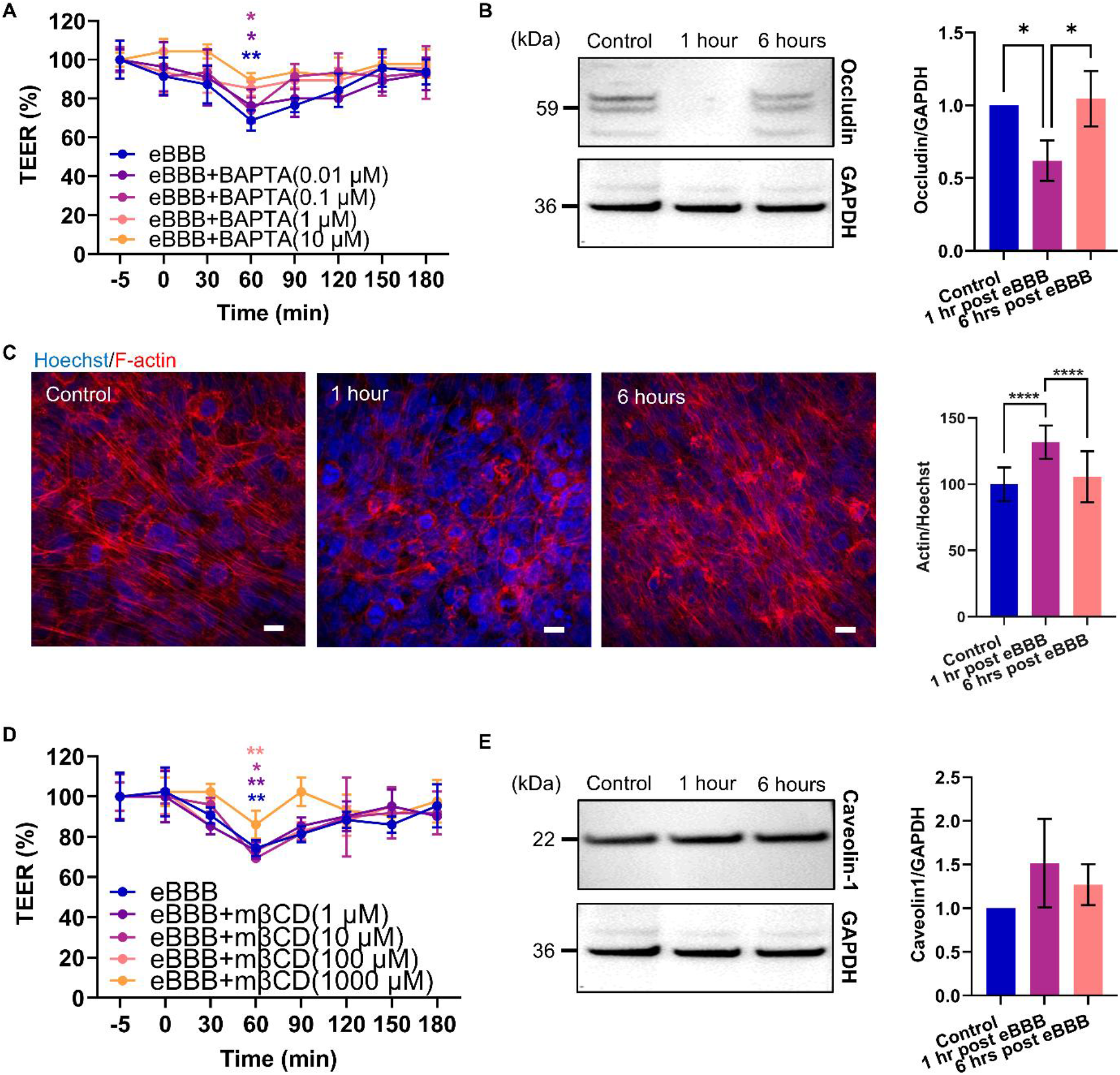
eBBB involves paracellular pathways. (A) Dose-dependent blockade of eBBB by BAPTA-AM. (B) Western blot analysis of occludin expression following eBBB treatment. (C) Immunofluorescence staining of F-actin showing acute cytoskeletal reorganization following eBBB. Scale bar = 10 µm. (D) Dose-dependent blockade of eBBB by mβCD. (E) Western blot analysis of caveolin-1 expression following eBBB treatment. n = 3 biological replicates. Statistical analysis: One-way ANOVA (*p < 0.05, **p < 0.01, ****p < 0.0001).

We next examined two downstream effectors of Ca²⁺ signaling at the endothelial junction: occludin, a transmembrane tight junction protein whose phosphorylation and internalization are regulated by Ca²⁺-activated kinases, and F-actin, whose polymerization into stress fibers drives paracellular gap formation (*37–39*). Occludin expression decreased to approximately 50% of baseline 1 hour after eBBB and recovered to control levels by 6 hours (**Fig. 4B**). In contrast, ES alone or stimulation in the presence of PLLA-PEG NPs did not significantly alter occludin expression (**fig. S4, A and B**), confirming that both nanoparticle-endothelial association and electrical activation are required to trigger junctional remodeling. Phalloidin staining showed an approximately 35% increase in F-actin signal at 1 hour, consistent with acute cytoskeletal reorganization into contractile stress fibers, followed by recovery to baseline at 6 hours (**Fig. 4C**).

The parallel kinetics of occludin downregulation, F-actin reorganization, and TEER reduction all peaking at 1 hour and recovering by 6 hours, consistent with a coordinated, Ca²⁺-mediated remodeling of the endothelial junction rather than nonspecific cytotoxicity.

We next investigated whether cholesterol-dependent membrane microdomains contribute to eBBB, because these domains support caveolar organization while also regulating the junctional localization and trafficking of endothelial TJ proteins (*40–42*). Pretreatment with methyl-β- cyclodextrin (mβCD), which depletes membrane cholesterol, dose-dependently attenuated the reduction in TEER, with the strongest inhibition observed at 1000 µM (**Fig. 4D**) (*43*). mβCD alone did not significantly alter TEER at the concentrations tested (**fig. S5**). In contrast, total caveolin-1 expression did not change significantly at either 1 or 6 hours after eBBB (**Fig. 4E**), nor following ES alone or stimulation in the presence of PLLA-PEG NPs (**fig. S4, C and D**). Because mβCD disrupts cholesterol-rich lipid rafts that anchor multiple TJ proteins, including occludin and ZO-1, in addition to serving as scaffolds for caveolae, its inhibitory effect on eBBB most likely reflects raft-dependent disruption of TJ protein anchoring rather than suppression of caveolar trafficking per se. The absence of a change in total caveolin-1 expression does not exclude a contribution from caveolin-1 phosphorylation-dependent vesicular trafficking, which regulates caveolar fission and cargo release independently of total protein levels (*44*).

Together, these findings define a Ca²⁺-dependent paracellular mechanism in which eBBB transiently activates cytoskeletal contraction, induces occludin internalization, and increases paracellular permeability, all of which fully reverse within 6 hours. The requirement for intact cholesterol-rich membrane domains, the absence of caveolin-1 upregulation, and the parallel recovery kinetics of junctional and cytoskeletal markers collectively support a predominantly paracellular route, while a concurrent transcellular contribution mediated by caveolin-1 phosphorylation cannot be excluded on the basis of total protein abundance alone.

### eBBB enables on-demand, localized, and reversible BBB modulation in the mouse brain

Having established that eBBB transiently increases endothelial permeability *in vitro*, we next evaluated its ability to modulate the BBB *in vivo* in mice. We employed a 4×1 HD-tDCS MEA consisting of a central anode surrounded by four symmetrically spaced cathodes. The electrodes were connected to a Soterix Medical transcranial electrical stimulator through a 4×1 HD-tES adaptor, which focuses current density within the targeted cortical region rather than distributing it across the entire brain, as occurs with conventional tDCS (**Fig. 5A**) To ensure consistent electrode positioning, the array was aligned using a prepatterned mold and embedded in polydimethylsiloxane (PDMS), preserving the geometry of the 4×1 montage (**Fig. 5B**). All electrodes were electrochemically coated with nano-Pt, which significantly reduced electrode impedance across the entire frequency spectrum (**Fig. S6**), indicating improved electrical conductivity, increased electrochemically active surface area, and enhanced charge transfer- capability, thereby enabling the safe delivery of HD-tDCS stimulation.

**Fig. 5.**
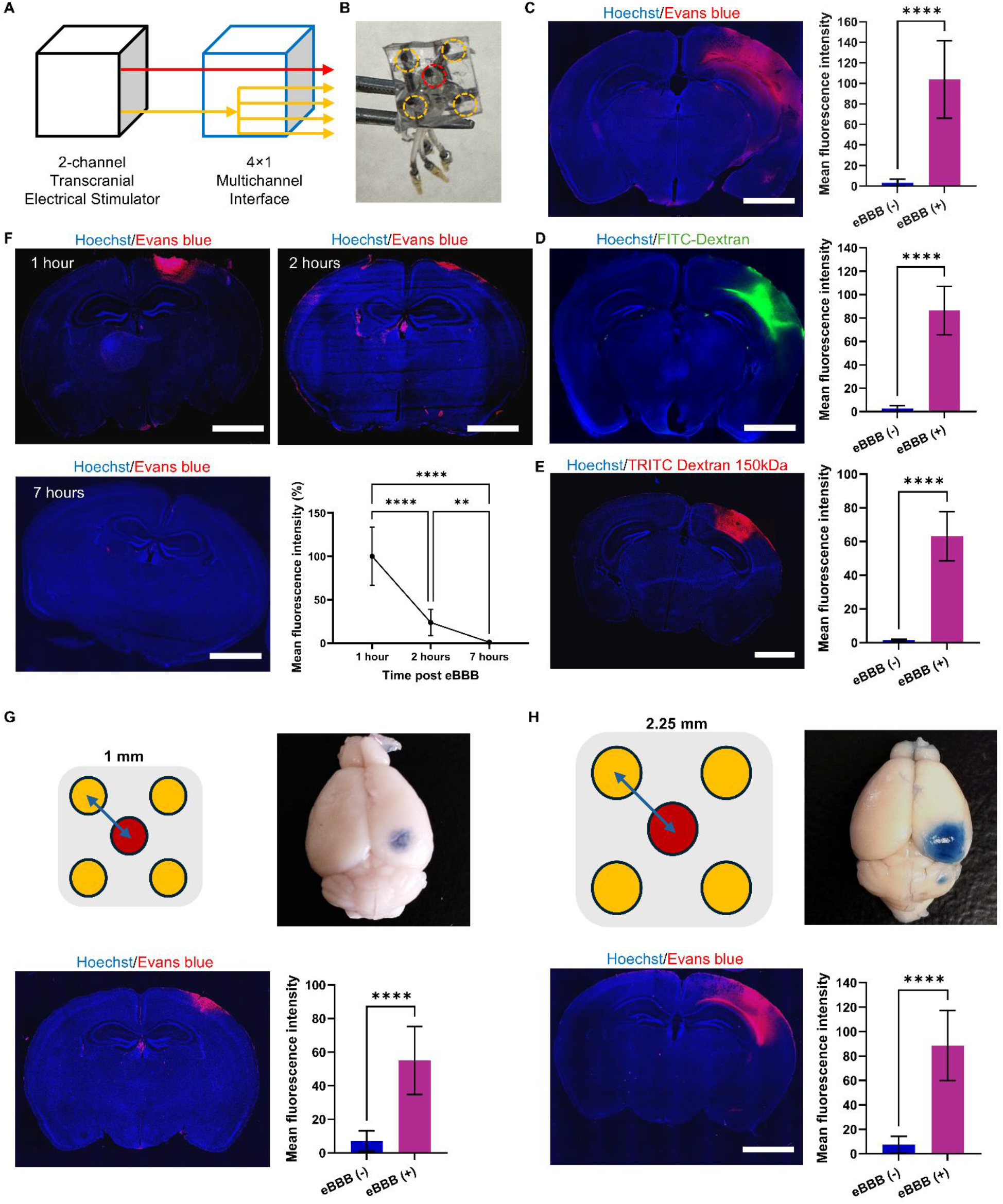
*In vivo* evaluation of eBBB. (A) Schematic of the 4×1 HD-tDCS concept. (B) Microelectrode array (MEA) configuration with one central anode surrounded by four cathodes. (C) eBBB increases the leakage of Evans blue (albumin-bound, 69 kDa) into the mouse brain. (D) eBBB increases the leakage of FITC-Dextran (70 kDa) into the mouse brain. (E) eBBB increases the leakage of TRITC-Dextran (150 kDa) into the mouse brain. (F) eBBB-induced BBB opening is reversible over time. (G-H) The spatial extent of BBB modulation can be tuned by varying electrode spacing on the MEA (4×1 HD-tDCS montage). In D–G, scale bar = 2 mm. n = 3 mice in C, F, G, and H. n>10 images from 1 mouse in D and E. Statistical analysis: Unpaired Student’s t-tests (C, D, E, G, H) or One-way ANOVA (F) (p < 0.01, **p < 0.0001).

We first examined how nanoparticle dose (40-150 mg/kg), stimulation current (0.2-0.8 mA), and stimulation duration (3-8 minutes) affected BBB permeability using Evans blue as a tracer. These studies identified 150 mg/kg PLLA-Alb NPs, 0.8 mA, and 8 minutes of stimulation as the conditions that produced the clearest localized response and were therefore selected for subsequent experiments (**fig. S7A-C**). Under these conditions, eBBB produced extravasation of 69-kDa Evans blue–bound albumin, 70-kDa FITC-dextran, and 150-kDa TRITC-dextran into the brain parenchyma (**Fig. 5C to E**). The progressive increase in tracer size from Evans blue–albumin to 150-kDa TRITC-dextran demonstrates that eBBB opens the paracellular pathway to macromolecular cargos substantially larger than the tight junction exclusion limit under normal conditions. Tracer accumulation was largely confined to the stimulated cortical region, with minimal leakage in adjacent tissue. Electrical stimulation alone, PLLA-Alb NPs without stimulation, and PLLA-PEG NPs combined with stimulation did not produce comparable Evans blue extravasation (**fig. S7D**), confirming that localized BBB modulation requires both endothelial-associated PLLA-Alb NPs and electrical activation. The spatial distribution of tracer leakage was also broadly consistent with the modeled electrical field beneath the electrode array generated using HD-Explore™ (**fig. S7E**).

We next evaluated the duration and spatial tunability of the response. BBB permeability was highest at 1 hour after eBBB, progressively returned toward baseline at 2 hours, and fully recovered within 7 hours, demonstrating that the opening was transient and reversible (**Fig. 5F**). Changing the distance between the central and surrounding electrodes from 1 to 2.25 mm altered the area of tracer extravasation, showing that the spatial extent of BBB opening can be adjusted through electrode geometry (**Fig. 5, G and H**), with the larger spacing producing a broader opening area consistent with the wider current density distribution. Together, these findings demonstrate that eBBB produces localized and reversible BBB modulation *in vivo*, with both the magnitude and spatial footprint of the permeability response controllable through treatment parameters and electrode configuration.

To determine whether this transient permeability increase was accompanied by lasting tissue injury, we examined major components of the NVU (**Fig. 6A**). 1 hour after eBBB, microvascular density remained unchanged, and the vascular abundance of ZO-1, claudin-5, and occludin did not differ significantly from controls at the time point examined (**Fig. 6, B to E**). Neuronal density (Nissl), as well as oligodendrocyte precursor cells and pericyte coverage (NG2), were also preserved (**Fig. 6, F to G**). The perivascular abundance of AQP4, normalized to CD31 signal, was similarly unchanged after eBBB (**Fig. 6H**), indicating that astrocytic end-feet polarization at the vascular interface was not disrupted. GFAP-positive astrocytes and Iba1-positive microglia increased transiently on day 3 but returned to baseline by day 7 (**fig. S8**), indicating a short-lived glial response rather than persistent neuroinflammation, consistent with the known reaction of brain tissue to localized mechanical or electrical perturbations that resolve without intervention (*23*). In a separate assessment of nanoparticle tolerability, mice receiving PLLA-Alb NPs maintained stable body weight and showed no overt histopathological abnormalities in major organs at the 13-week long-term endpoint (**fig. S9**). These data support the systemic biocompatibility of PLLA-Alb NPs over a clinically relevant observation window. Together, these findings show that eBBB creates a localized, tunable, and reversible permeability window without lasting disruption of neurovascular structure at the time points examined, supporting its use for regional brain delivery.

**Fig. 6.**
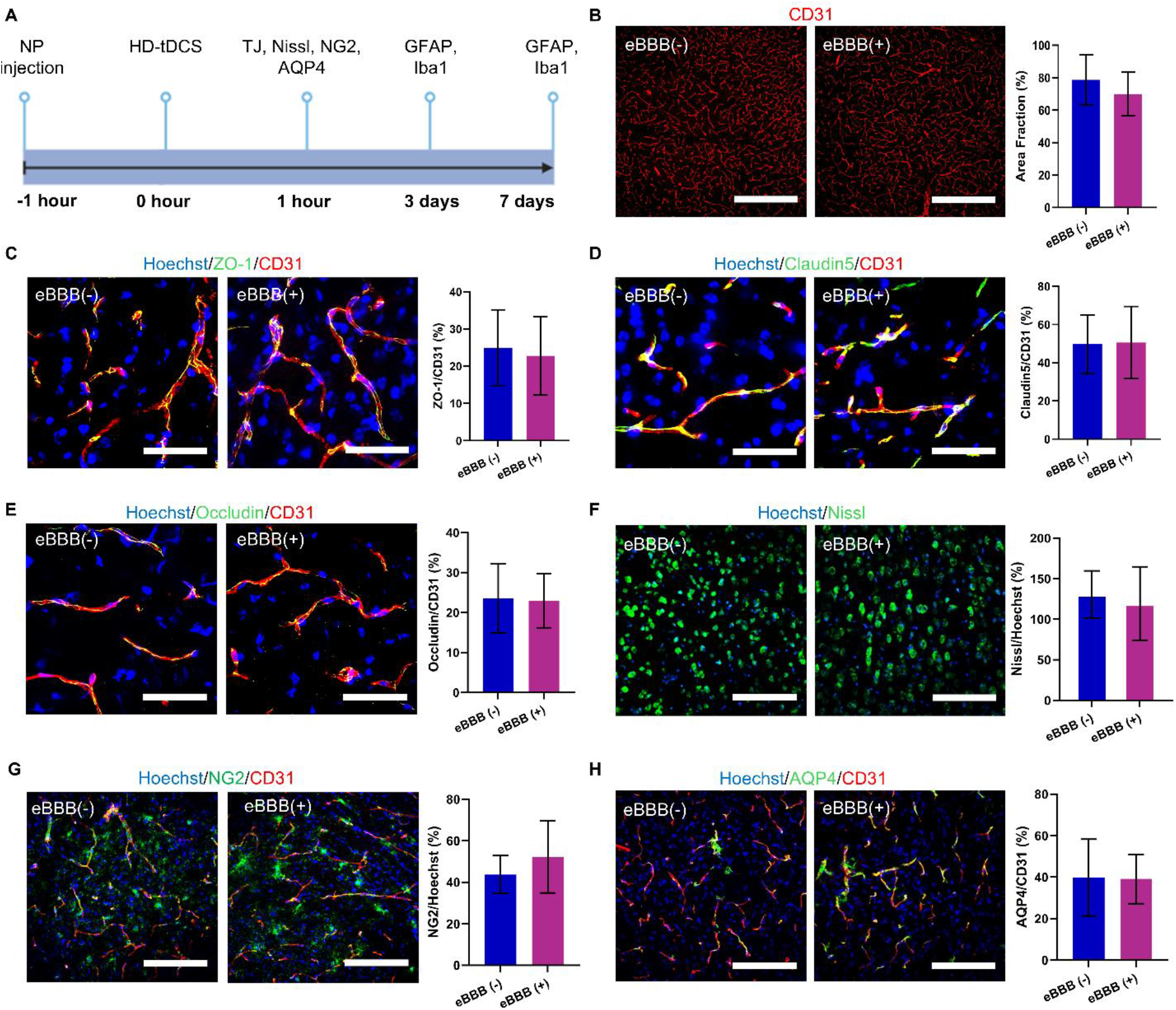
eBBB does not induce lasting disruption of neurovascular unit architecture. (A) Experimental timeline showing the schedule of eBBB treatment and tissue collection for immunohistochemical staining of neurovascular unit components at the indicated time points. (B) Representative images and quantification of CD31⁺ microvascular density in the eBBB-treated cortex versus contralateral side. (C) Representative images and quantification of ZO-1 expression. (D) Representative images and quantification of claudin-5 expression. (E) Representative images and quantification of occludin expression. (F) Representative images and quantification of Nissl staining. (G) Representative images and quantification of NG2 coverage. (H) Representative images and quantification of AQP4 expression. For panels B to H, data are presented as mean ± SD. n = 3 mice per group. Statistical analysis: Unpaired Student’s t-tests. ns, not significant (p > 0.05) for all comparisons. Scale bars = 400 µm in (B) and 50 µm in (C-H).

### eBBB enhances regional delivery of diverse therapeutic cargos

To determine whether the localized permeability window created by eBBB could support therapeutic delivery, we evaluated three cargo classes that are normally excluded by the intact BBB and represent distinct modalities of CNS treatment: a small-molecule drug, a full-length antibody, and an AAV gene therapy vector. We first examined Taxol646, a far-red fluorescent paclitaxel analog that retains the microtubule-stabilizing activity of the parent compound and has been used to track taxane distribution in tissue (*14*). Following systemic administration, Taxol646 accumulation in the eBBB-treated brain region was 33-fold higher than in mice that received the drug without eBBB, with the fluorescence concentrated within the stimulated cortex (**Fig. 7A**), confirming regional specificity. These findings demonstrate that eBBB can enhance parenchymal accumulation of a clinically relevant anticancer agent in a spatially defined cortical region, which is directly applicable to the treatment of superficial brain metastases and frontocortical gliomas. We next assessed whether eBBB could facilitate delivery of a full-length antibody, a cargo class of growing therapeutic relevance given the increasing number of CNS-targeting immunotherapies and checkpoint inhibitors under clinical investigation (*45, 46*). Following systemic administration of rabbit IgG, fluorescence intensity in the eBBB-treated hemisphere was 15-fold higher than in the contralateral hemisphere **(Fig. 7B)**. Parenchymal IgG signal was distributed within the stimulated cortical region, consistent with the spatial footprint of BBB opening established by tracer studies.

**Fig. 7.**
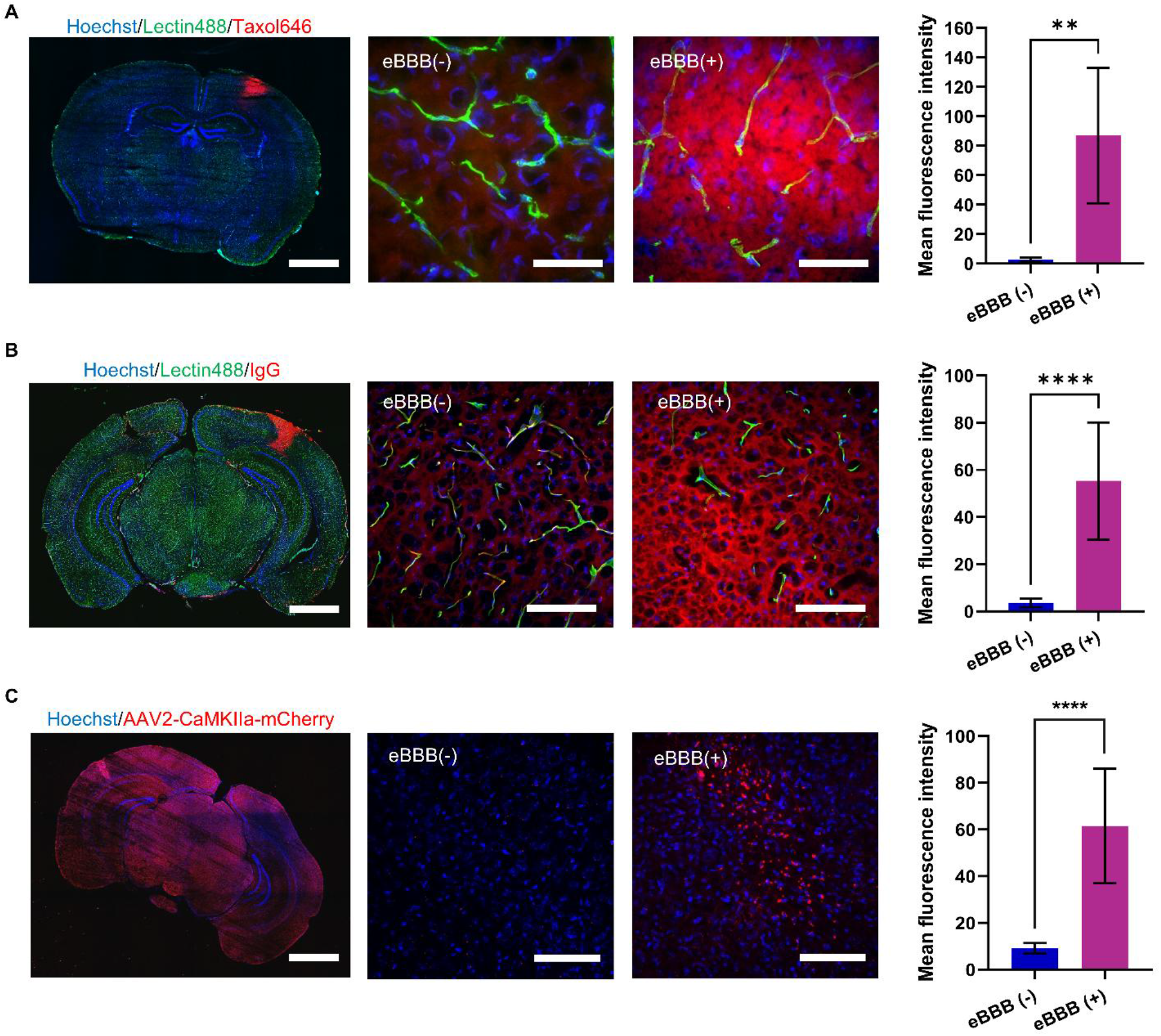
eBBB enhances regional brain delivery of pharmacologically diverse therapeutic cargos. (A) Regional delivery of a small-molecule anticancer drug, Taxol646, and quantification of mean fluorescence intensity in the eBBB (-) and eBBB (+) regions. (B) Regional delivery of a full-length antibody, rabbit IgG, and quantification of mean fluorescence intensity in the eBBB (-) and eBBB (+) regions. (C) Regional delivery of an adeno-associated viral gene therapy vector, AAV2-CaMKIIa-mCherry, and quantification of mean fluorescence intensity in the eBBB (-) and eBBB (+) regions. For all panels, data are presented as mean ± SD. n>10 images per group. Statistical analysis: Unpaired Student’s t-tests; **p < 0.01, ****p < 0.0001. Scale bar = 2 mm in the low-magnification images and 100 µm in the high-magnification images.

We next examined whether eBBB could facilitate delivery of an AAV, a widely used vehicle for *in vivo* gene therapy. Following systemic administration of AAV2-CaMKIIα-mCherry (5×10¹¹ vg per mouse, intravenous), mCherry fluorescence intensity in the eBBB-treated hemisphere was quantified at 4 weeks post-injection, a timeframe sufficient for stable transgene expression under the CaMKIIα promoter. Regional mCherry fluorescence was 7-fold higher in the eBBB-treated cortex than in the contralateral hemisphere (**Fig. 7C**), demonstrating that eBBB-enhanced AAV2 entry led to increased neuronal transduction in the targeted region.

Together, these findings demonstrate that eBBB enhances regional brain delivery across three pharmacologically and mechanistically distinct cargo classes. The ability to deliver such diverse cargos through a single, electronically controlled BBB-opening platform substantially broadens the translational scope of eBBB and supports its application to CNS malignancies and neurological disorders.

## Discussion

The eBBB platform introduced here addresses a fundamental challenge in CNS drug delivery by enabling on-demand, spatially localized, and reversible modulation of the BBB. Through HD-tDCS-mediated activation of vascular-anchored PLLA-Alb NPs, BBB permeability can be controlled with high spatial and temporal precision, making this approach particularly well suited for cortical and other superficial brain regions. To the best of our knowledge, this is the first demonstration of HD-tDCS-driven activation of piezoelectric nanoparticles to modulate BBB permeability, establishing a mechanistically distinct paradigm within the broader landscape of physical BBB opening strategies.

The translational potential of eBBB is supported by two well-established regulatory precedents. First, PLLA is an FDA-approved biodegradable polymer that has been widely used in implantable medical devices and tissue-engineering scaffolds, providing a well-characterized safety foundation for PLLA-based biomaterials (*47*). Second, HD-tDCS has been evaluated in clinical trials across diverse neurological and psychiatric disorders, demonstrating a favorable safety profile (*48–50*). Together, these precedents support the clinical feasibility of the eBBB platform. Moreover, its compact, low-infrastructure design reduces barriers to deployment in both research and clinical environments, making eBBB well suited for broad implementation and future translation.

The mechanistic findings reported here extend our understanding of how mechanical forces at the endothelial surface regulate BBB permeability. We previously demonstrated that mechanical stimulation of brain ECs elicits transient Ca²⁺ elevation, actin polymerization, and ERK1/2 phosphorylation, leading to increased paracellular permeability (*24*). The present study confirms that eBBB activates a similar Ca²⁺-dependent signaling cascade, as evidenced by the dose-dependent inhibition of barrier opening by the intracellular Ca²⁺ chelator BAPTA-AM. This response is accompanied by transient occludin downregulation and F-actin reorganization, both of which return to baseline within 6 hours, consistent with the reversible nature of BBB modulation. Although total caveolin-1 expression was unchanged, caveolae-mediated transcytosis is regulated by caveolin-1 phosphorylation and the formation, fission, and trafficking of caveolar vesicles (*44, 51*). Thus, the present findings do not exclude a concurrent transcellular contribution to eBBB- induced permeability.

The spatial precision of eBBB is achieved through two complementary mechanisms: the HD-tDCS electrode montage confines the electrical field to the targeted cortical region, and BSA-functionalized PLLA NPs promote the association with the brain microvasculature, ensuring that only NPs within the stimulated volume are activated. Spatial confinement is further enabled by our custom-designed nano-Pt-coated MEAs, whose nanostructured morphology increases electrode surface area and reduces impedance, ensuring stable current injections without fluctuations that could compromise safety or efficacy. Embedded in PDMS substrates, these MEAs conform seamlessly to skull contours for consistent transcranial delivery. Importantly, the extent of BBB opening can be tuned by adjusting electrode spacing and stimulation parameters, providing flexibility to target applications ranging from focal lesions to larger cortical areas.

ZO-1 downregulation was observed *in vitro* by Western blot but was not detected by IHC *in vivo* when TJ protein expression was normalized to the CD31 signal. Several factors are likely to contribute to this discrepancy. The hCMEC/D3 monoculture lacks the paracrine support of pericytes and astrocytes, which actively maintain TJ protein expression in the intact neurovascular unit and may accelerate recovery beyond the sensitivity of IHC detection at the time points examined (*52, 53*). Moreover, hCMEC/D3 cells exhibit reduced expression of key BBB junctional genes, including *CLDN5* and *OCLN*, relative to freshly isolated brain endothelial cells, rendering this model more sensitive to transient junctional remodeling than the intact BBB (*54–56*).

Several limitations of the present study should be acknowledged. First, although the current data establish that eBBB requires both PLLA-Alb nanoparticles and electrical stimulation, future studies can further define its electromechanical basis by quantifying the piezoelectric response of the nanoparticles and correlating this response with BBB permeability. Second, the *in vitro* BBB model employed lacks the full complexity of the neurovascular unit, including pericytes, astrocytes, and fluid shear stress; more physiologically complete models, such as microfluidic BBB-on-a-chip systems or organotypic brain slice preparations, or *in vivo* models, should be used to further validate the BBB opening mechanism. Third, all *in vivo* experiments were performed in healthy mice with an intact BBB; the behavior of PLLA-Alb NPs and the efficiency of eBBB- induced opening may differ in the context of neurological disease, where the BBB is already partially compromised, and vascular architecture is altered. Moreover, while body weight stability and histological analysis after a single eBBB modulation demonstrated no overt toxicity, longer-term safety studies, including repeated dosing over weeks to months, will be required to support clinical translation. Lastly, the spatial resolution of BBB opening, while tunable by electrode montage, is ultimately constrained by the physics of transcranial current spread; achieving sub-millimeter precision in humans, where the skull is thicker and the brain larger than in mice, will require computational modeling and individualized electrode optimization.

eBBB establishes a new class of BBB modulation that couples nanomaterial-mediated vascular targeting with programmable transcranial electrical stimulation to achieve on-demand, spatially controlled, and reversible CNS access. The platform’s key attributes- component-dependent activation, tunable spatial footprint, multimodal cargo delivery, and absence of lasting neurovascular damage- collectively address unmet needs in regional brain drug delivery. Looking forward, eBBB is particularly well-suited for superficial CNS targets accessible to transcranial stimulation, including frontocortical gliomas, brain metastases, and neurodegenerative foci, and its modular design, separable nanoparticle and stimulation components, enables independent optimization of vascular targeting and spatial field patterning as the platform advances toward clinical translation.

## Materials and Methods

### Chemicals and supplies

Poly(L-lactide) acid endcap (Mn 45,000-55,000 Da) was purchased from Akina Inc. Bovine serum albumin, Molecular Probes™ Tetramethylrhodamine-5-(and-6)-Isothiocyanate (5(6)-TRITC) mixed isomers, Dichloromethane, Ethylacetate, Dimethyl formamide, Dimethyl sulfoxide, Oxalyl chloride, Trehalose dihydrate, Invitrogen™ BAPTA-AM (cell permeant chelator), Chlorpromazine hydrochloride (>98%), Methyl-β-cyclodextrin (>98%), Invitrogen™ Alexa Fluor™ 647 Phalloidin, Triton-X 100, SouthernBiotech™ normal donkey serum, goat serum, goat anti-CD31 (AF3628), rabbit anti-Claudin-5 (34–1600), rabbit anti-ZO-1(40–2200), rabbit anti-VE-cadherin (36–1900), rabbit anti-Occludin (71-1500), rabbit anti-NG2 (AB5320), rabbit anti-GFAP (PA1-10019), rabbit anti-Iba1 (AB178846), mouse anti-AQP4 (sc-32739), rabbit IgG isotype control (02-6102), NeuroTrace 500/525 (N21480), donkey anti-goat IgG Alexa 488 (A-11055), donkey anti-rat IgG Alexa 594 (A-21209), donkey anti-rabbit IgG Alexa 594/647 (A-21207, A- 31573), Thermo Scientific™ RIPA Lysis and Extraction Buffer, Thermo Scientific™ Halt™ Protease Inhibitor Cocktail (100X), Thermo Scientific™ Halt™ Phosphatase Inhibitor Single-Use Cocktail, Thermo Scientific™ Pierce™ Bovine Serum Albumin Standard, 2 mg/mL, Bradford Protein Assay Kit, Invitrogen™ NuPAGE™ Transfer Buffer (20X), Invitrogen™ NuPAGE™ MES SDS Running Buffer (20X), Thermo Scientific™ SuperSignal™ West Pico PLUS Chemiluminescent Substrate, Invitrogen™ NuPAGE™ Bis-Tris Mini Protein Gels (4-12%, 1.0- 1.5 mm), GAPDH Polyclonal Antibody, MilliporeSigma™ Immobilon™-PSQ PVDF Membranes, Invitrogen™ SDS Protein Sample Loading Buffer, and Thermo Scientific™ PageRuler™ Plus Prestained Protein Ladder (10 to 250 kDa), cell culture grade endotoxin-free water, sterile 10x PBS, FITC-dextran (40 kDa, 70 kDa), TRITC-Dextran (150 kDa), FGF-2, trypsin-EDTA, collagen type I, DMSO, sucrose, Penicillin–streptomycin, trypan blue, Cayman Chemical WST-1 Cell Proliferation Assay Kit, Hoechst dye 33342, LIVE/DEAD™ Cell Imaging Kit (488/570), Evans blue dye, sterile saline, paraformaldehyde, lectin Dylight 488/594/647, Tissue-Plus™ O.C.T. compound, and Invitrogen Fluoromount-G™ mounting medium, and Spectra/Por^®^ Dialysis Membrane, MWCO: 300 kDa were purchased from Fisher Scientific. Rat anti-CD31 (550274) was purchased from BD Biosciences. Amine-PEG-Carboxymethyl (MW 5,000) was purchased from Laysan Bio Inc. Caveolin-1 (D46G3) XP® Rabbit mAb was purchased from Cell Signaling Technology. All chemicals were analytical grade unless specified. 12 and 96- well plates, and Falcon® cell culture inserts for 12-well plates with 0.4 µm transparent PET membrane inserts were purchased from VWR. Millicell ERS-PEG 50002 System was purchased from Millipore. EVOM2 cell culture cuvette chamber for TEER measurement was purchased from World Precision Instruments. 1x1 transcranial Electrical Stimulation Device and 4×1 HD-tDCS/HD-tES Adaptor were purchased from Soterix Medical.

### Cell culture

Human cerebral microvascular endothelial cells (D3 cell line) and the EndoGRO™-MV Complete Media Kit were purchased from Millipore.

### Animals

All animal procedures were performed in accordance with the Guidelines for Care and Use of Laboratory Animals of Louisiana State University and approved by the Institutional Animal Care and Use Committee (IACUC). Adult female C57BL/6 mice (7 weeks old, 20-25 g) were ordered from the Jackson Laboratory. Potential sex-dependent effects were not evaluated in this study. All animals were bred in pathogen-free conditions, in temperature (20-22 °C) and humidity (52-57%)- controlled housing, under a 12-h light/dark cycle, and with free access to food and water. Mouse sex was not considered in the study design.

### Fabrication of electrodes

For *in vitro* experiments, the electrodes were fabricated using tungsten wires (6 cm in length, 508 µm in diameter) due to their high stiffness, which facilitated precise handling and positioning. The wires were insulated using a thin layer of medical-grade epoxy (AA-BOND FDA16 Medical Grade Epoxy Adhesive), leaving a 4 mm segment at the tip intentionally exposed for stimulation. To improve surface roughness and enhance nano-platinum (nano-Pt) adhesion, the exposed tungsten was electrochemically etched in a 2 M KOH solution at 10 V for 100 seconds. Following etching, the exposed tungsten was coated with nano-Pt as described below.

For *in vivo* experiments, we custom-designed stimulation devices with a 4-by-1 electrode montage, consisting of a central anode surrounded by four cathodes. The cathodes were symmetrically spaced 2.25 mm from the central anode. The five electrodes were constructed from the cross-section of insulated silver wires (650 µm diameter), each coated with nano-Pt. To precisely align the five electrodes in the desired montage configuration, we used a pre-patterned SU-8 mold with holes sized to accommodate the wires. The electrode assembly was subsequently encapsulated in polydimethylsiloxane (PDMS) to maintain structural integrity.

### SU-8 mold fabrication

A ∼10 µm thick SU-8 layer was spin-coated at 2000 rpm onto a SiO₂ wafer and patterned via standard negative photolithography (exposure dose: 250 mJ/cm²). After development, the SU-8 molds were released from the SiO₂ substrate using buffered oxide etchant. Anchor holes in the mold allowed precise positioning of the silver wires, which were then encapsulated in PDMS to form the final electrode array.

### Nano-platinum coating

All microelectrodes were electrochemically coated with nano-Pt by cycling the potential between -0.35 V and 0.3 V vs Ag/AgCl in an aqueous solution containing 1 mM HCl and 25 mM H₂PtCl₆.

The deposition was carried out for 300 cycles to ensure consistent coating across all electrodes.

### Electrochemical characterization

Electrochemical impedance spectroscopy (EIS) and cyclic voltammetry (CV) were used to measure impedance spectra and charge storage capacity before and after the nano-Pt coating and verify the capability of the electrodes to sustain the required stimulation (1 mA for 5 minutes). EIS and CV were performed in 1×PBS in a three-electrode electrochemical cell set-up with a platinum counter electrode and an Ag/AgCl wire reference electrode, using a potentiostat/galvanostat (Autolab, Metrohm, Riverview, FL, USA). During the EIS measurements, a sine wave (10 mV RMS amplitude) was superimposed onto the open-circuit potential while varying the frequency from 1 to 105 Hz. During the CV tests, the working-electrode potential was swept between 0.8 and -0.6 V vs. Ag/AgCl, maintaining a scan rate of 100 mV/s.

### Stability test

To assess the stability of the nano-Pt coated electrodes under stimulation conditions, we applied a constant current of 1 mA (the highest used) for 5, 10, and 20 minutes. During stimulation, the electrode potential was continuously monitored using a two-electrode configuration with a reference platinum electrode. The test was conducted in 1x PBS. Pre- and post-stimulation EIS and CV were recorded to evaluate any degradation in electrode performance. Visual inspection under a microscope was also performed to identify potential delamination or structural damage to the nano-Pt coating.

### Synthesis of PLLA conjugates

PLLA–albumin (PLLA–Alb), PLLA–PEG, and PLLA–TRITC were prepared using a common oxalyl chloride/DMF activation and coupling procedure. PLLA was dissolved in dichloromethane (DCM), activated with oxalyl chloride and dimethylformamide (DMF) for 5 hours at room temperature, and subsequently reacted overnight with albumin or TRITC dissolved in anhydrous dimethyl sulfoxide (DMSO), or with Amino-poly(ethylene glycol) carboxy methyl ether (NH₂– PEG–CM) dissolved in DCM. TRITC-containing reactions were protected from light. The resulting conjugates were precipitated with diethyl ether, redissolved in DCM, washed four times with ultrapure water, concentrated by rotary evaporation, and lyophilized for 48 hours. Formulation-specific reagent amounts are summarized in **Table S1**.

### Preparation of PLLA nanoparticles

Unlabeled and TRITC-labeled PLLA–Alb and PLLA–PEG NPs were prepared by oil-in-water emulsion–solvent evaporation using the conditions provided in **Table S2**. The appropriate PLLA conjugate or conjugate blend was dissolved in the organic phase and emulsified in ultrapure water. Next, the mixture was probe-sonicated for 30 minutes using the 4-seconds ON, 2-second OFF pulse setting at 50% amplitude to produce an oil-in-water emulsion (Sonics,Vibra cell TM, Model CV33). Fluorescent nanoparticles were prepared by incorporating PLLA–TRITC into the corresponding PLLA–Alb or PLLA–PEG polymer feed before emulsification. Ethyl acetate was included in the PLLA–PEG/TRITC formulation to dissolve PLLA–TRITC. Organic solvents were removed by rotary evaporation at 33 °C under reduced pressure for 2 hours. Trehalose and PVA were used in the Alb- and PEG-based formulations, respectively. The resulting nanoparticles were lyophilized for 48 hours and stored at −20 °C until use.

### Nanoparticle characterization

The nanoparticles were characterized by Zeta View® Particle Metrix PMX120 for hydrodynamic size and zeta potential, and Transmission Electron Microscopy (TEM) (JEOL JEM 1400, Peabody, MA) for size and shape. Before use, the NPs were dialyzed using a 300 kDa MWCO dialysis membrane in 1× PBS for 24 hours.

### Protein quantification

Protein concentration was determined using the Bradford Assay based on the binding of Coomassie dye to proteins. Briefly, 5 µL of PLLA-Alb NP solution after the purification step was mixed with 250 µL of Bradford reagent and incubated at room temperature for 5–10 minutes. Absorbance was measured at 595 nm using a microplate reader/spectrophotometer. A standard calibration curve was generated using bovine serum albumin (BSA) standards of known concentrations: 2000, 1500, 1000, 750, 500, 250, 125, and 0 µg/mL. Protein concentrations of the nanoparticle were calculated from the standard curve and expressed as mg/mL.

### Cell culture and the *in vitro* BBB model

Human brain microvascular endothelial cells (hCMEC/D3 cells, passage <10) were cultured in EndoGRO™-MV Complete media supplemented with FGF-2 (1 ng/mL). The transwell inserts (0.4 µm pore size, 0.9 cm^2^) were coated with Collagen type I (15 µg/mL, 500 µL) for 1 hour at 37 °C. After rinsing with PBS, the hCMEC/D3 cells were seeded in transwell inserts at a density of 90,000 cells/cm² in 0.6 mL of cell culture medium, placed in a 12-well plate filled with 1 mL of cell culture medium, and cultured for 7 days at 37 °C and 5% CO₂ to form cellular monolayers.

The integrity of the hCMEC/D3 monolayers was characterized by transendothelial electrical resistance (TEER) and permeability. The expression of tight junction protein was evaluated using immunocytochemistry staining and Western Blot.

### Transendothelial electrical resistance (TEER) measurement

The Transwell insert was placed in the Cell Culture Cup Chamber (WPI) connected to the epithelial voltmeter (Millicell ERS-2) for TEER measurement, following the manufacturer’s instructions under room temperature. The resistance value of blank culture inserts coated with collagen on the top side of the membrane was used as the baseline. To obtain the TEER, the baseline value (R_blank_) was subtracted from the resistance measured from the inserts cultured with cells (R_total_). The resulting resistance value, multiplied by the effective membrane area (A, cm²), yields the TEER value in Ω cm².

TEER = (R_total_- R_blank_) × A

### Permeability measurement

0.5 mL of FITC-dextran (40 kDa, 1 mg/mL in medium) was added to the upper compartments of the Transwell inserts. At particular time points, i.e., 0, 30, 60, 90, 120, 150, 180, 210, and 240 minutes, 100 μL of the medium was aspirated from the bottom well and added to 96-well plates. 100 μL of medium was then replaced in the bottom chamber to keep the volume constant. We then measured the fluorescent intensity for the collected samples in the 96-well plate (excitation at 490 nm and emission at 540 nm) to obtain the FITC-dextran concentration. The quantity (*Q*) was obtained by multiplying the concentration and volume. The apparent permeability (P_app_, cm s^−1^) was calculated by the rate of FITC-dextran quantity change over time (dQ/dt, mg/s), divided by the initial concentration of dextran (C₀, 1 mg/mL) and the membrane area (A, cm^2^).

P_app_ = (dQ/dt) / (A × C₀)

### Nanoparticle Uptake

0.5 mg/mL of PLLA-Alb-TRITC or PLLA-PEG-TRITC NPs were incubated with the hCMEC/D3 cell monolayer for 30 minutes, followed by three rinses with 1×PBS. Then, the cells were fixed with 4% PFA for 15 minutes at 4 °C. After rinsing with 1×PBS, the cell nuclei were stained with Hoechst for 10 minutes. Finally, the cells were washed three times with 1×PBS before imaging.

### Cell Viability

hCMEC/D3 cells (3000 cells) in 100 µL of cell culture medium were seeded in each well of the 96-well plate. A day later, PLLA-Alb NPs with a variety of concentrations (i.e., 0, 0.01, 0.1, 0.5, and 1 mg/mL) were added to each well, and the solution was gently mixed and incubated in a CO_2_ incubator (37 °C) for 72 hours. Then, cell viability was measured using the WST-1 assay, with absorbance readings taken at 450 nm according to the manufacturer’s instructions. 6 replicates were performed for each concentration.

### *In vitro* BBB modulation

The cells were incubated with PLLA-Alb NPs at 0.01, 0.1, 0.5, or 1 mg/mL for 30 minutes at 37 °C and 5% CO_2_. The treated cellular monolayer was washed three times with PBS. Two nano-Pt electrodes were positioned on opposite sides of the inserts, with 4 mm of each electrode tip submerged in the cell culture medium. The electrodes were connected to the Soterix Medical animal tES device. Then, the cells were stimulated with a variety of currents (i.e., 0.02 mA, 0.05 mA, or 0.1 mA for 1, 2, or 5 minutes (including a 30-second ramp-up and 30-second ramp-down). The *in vitro* BBB opening efficiency and safety were characterized by TEER and permeability measurements.

### *In vitro* calcium chelation by BAPTA-AM

To evaluate the role of intracellular calcium signaling in PLLA-Alb nanoparticle-mediated eBBB modulation, BAPTA-AM was used as a calcium chelator. A 10 mM BAPTA-AM stock solution was prepared in DMSO (3.82 mg in 500 μL), and working solutions (10, 1, 0.1, and 0.01 μM) were prepared by serial dilution in complete culture medium. hCMEC/D3 BBB models were first treated with PLLA-Alb NP (0.5 mg/mL) for 30 minutes, followed by washing with 1x PBS to remove unbound nanoparticles and TEER measurement. BAPTA-AM (500 μL/insert) was then applied for 30 minutes, followed by washing with 1x PBS and TEER measurement prior to electrical stimulation. The BBB models were stimulated at 0.1 mA for 5 minutes, and TEER was monitored immediately after stimulation (0 minutes) and at 30, 60, 90, 120, 150, and 180 minutes. PLLA- Alb nanoparticle-treated models without BAPTA-AM treatment were used as controls.

### *In vitro* caveolae inhibition by methyl β-cyclodextrin (MβCD)

To evaluate the role of caveolae-mediated transport in PLLA-Alb nanoparticle-induced eBBB modulation, MβCD was used as a caveolae-disrupting agent. A 10.48 mM MβCD stock solution was prepared in PBS (20.6 mg in 1.5 mL), and working concentrations (1, 10, 100, and 1000 μM) were prepared by dilution in complete culture medium. BBB models were first treated with PLLA- Alb NPs (0.5 mg/mL) for 30 minutes, followed by washing with 1x PBS and TEER measurement. MβCD (500 μL/insert) was then applied for 60 minutes, followed by washing with 1x PBS and TEER measurement prior to electrical stimulation. Samples were stimulated at 0.1 mA for 5 minutes, and TEER was monitored immediately after stimulation (0 minutes) and at 30, 60, 90, 120, 150, and 180 minutes. PLLA-Alb nanoparticle-treated models without MβCD treatment were used as controls.

### Live and dead cell staining

1 hour after eBBB, the cell viability was performed using live/dead cell staining according to the manufacturer’s instructions.

### Western blot

D3 cell monolayers were incubated with PLLA-Alb NPs (0.5 mg/mL) for 30 minutes, followed by rinsing with 1xPBS and electrical stimulation. The total protein extraction was performed using RIPA for 30 minutes, followed by rinsing with 1xPBS and electrical stimulation. The total protein extraction was performed using RIPA buffer at various time points, following the manufacturer’s instructions. Briefly, the cells were washed twice with cold PBS, followed by incubation in cold RIPA buffer mixed with a protease inhibitor and a phosphatase inhibitor cocktail for 5 minutes on ice with gentle swirling. The lysate was centrifuged (14000 g, 15 minutes) to collect the supernatant. Total protein concentration was determined using the Bradford assay kit. The exact amounts of protein (20 μg) were loaded onto a stain-free protein gel and separated by electrophoresis at 150 V. The proteins were then transferred to the LF PVDF membrane with the trans-Blot Turbo system (Bio-Rad), followed by blocking with 3% BSA in TBS buffer at room temperature. After 1 hour, the membranes were incubated with primary antibodies at 4 °C overnight. After washing the primary antibodies with TTBS buffer, the corresponding secondary antibodies were applied at room temperature for 1 hour. The blots were enhanced with chemiluminescence and then visualized using a ChemiDoc Touch Imaging System (Bio-Rad). The band intensity was analyzed using Fiji/ImageJ.

### Immunocytochemistry staining

For occludin staining, the monolayer was fixed using 4% PFA and incubated in blocking solution (5% Normal Donkey Serum + 0.1% Triton X-100) for 1 hour at room temperature. After washing 3 times with PBS, the cells were treated with the Occludin primary antibody (1:500 in blocking buffer) overnight at 4 °C. After rinsing 3 times with PBS, the cells were incubated with the secondary antibody (1:500 in PBS) and Hoechst dye (1:2000 in PBS) for an hour. After rinsing 3 times with PBS, the membrane of the insert was then cut and mounted on glass slides for fluorescent imaging.

For F-actin staining, the monolayer was fixed with 4% PFA for 15 minutes at 4 °C. After 3 washes with PBS, the cells were incubated with phalloidin-647 (1:1000) at room temperature for 1 hour, followed by Hoechst staining (1:2000). After rinsing 3 times with PBS, the membrane of the insert was then cut and mounted on glass slides for fluorescent imaging.

### *In vivo* BBB opening

The mice were anesthetized by 2-3% isoflurane (in air) and the PLLA-Alb or PLLA-PEG was intravenously injected. 1 hour later, the mouse was placed on a stereotactic frame with the skin on its head gently opened. The multielectrode array was mounted on the stereotaxic arm and connected to the Soterix stimulator. The multielectrode array was positioned conformally over the skull to cover the target brain region. A thin layer of high-definition electrolyte gel (Soterix Medical) was applied to the skull, acting as the buffer between the skull and the electrodes. To optimize BBB modulation and achieve the highest opening efficiency, different nanoparticle doses (40-150 mg/kg, 100 µL), current (0.2-0.8 mA), and stimulation times (3-8 minutes) were tested. Evans blue dye (2% in PBS, 100 µL) was injected intravenously to visualize BBB modulation. To determine the BBB recovery, Evans blue dye was injected at 1 hour, 2 hours, and 7 hours after BBB modulation. 30 minutes later, the mice were perfused with ice-cold PBS and 4% PFA, and the brains were extracted and post-fixed with 4% PFA overnight. The brains were frozen on dry ice and cut into 20 µm-thick slices using a cryostat. The slices were stained with Hoechst dye (1:2000) before imaging.

### Cargo delivery studies

For Taxol646 delivery, 1 mg of the drug was dissolved in 50 µL of DMSO to make 20mg/ml stock solution and diluted in saline to achieve a final concentration of (2.5mg/mL). Lectin 488 (100 μL) and diluted Taxol646 solution (100 μL) were i.v. injected into a mouse after eBBB. After 1 hour, the animal was perfused with ice-cold PBS and 4% PFA. For antibody delivery, lectin 488 (100 μL) and rabbit IgG (5 mg/mL) were i.v. injected into a mouse after eBBB. After 1 hour, the animal was perfused with ice-cold PBS and 4% PFA. For AAV delivery, AAV2-CaMKIIa-mCherry (5×10^11^ vg/mL, 100 μL) was injected into the tail vein of a mouse treated with eBBB. After 4 weeks, the animal was perfused with ice-cold PBS and 4% PFA.

All brain tissues were post-fixed with 4% PFA at 4 °C overnight. The brain tissue was then dehydrated in 30% sucrose, sliced to 30 μm thickness, and stained with Hoechst. Rabbit IgG delivery was assessed by staining the tissue with an anti-rabbit secondary antibody.

### Immunohistochemistry staining

To immunostain the vascular biomarker CD31, BBB tight junction proteins (Claudin-5, ZO-1, Occludin), and pericyte (NG2), the mouse brain tissue was snap-frozen on dry ice immediately after removal from the skull, stored at -20 °C, and then processed at 30 μm on a cryostat. The coronal brain sections were fixed for 10 minutes using ice-cold methanol at -20 °C. Blocking solution (5% normal donkey serum, 0.1% Triton X-100 in PBS) was applied to the tissue for 1 hour at room temperature. After washing, the sections were incubated overnight at 4 °C with primary antibodies (diluted 1:500). After washing, the sections were incubated for 1 hour at room temperature with secondary antibodies, followed by incubation with Hoechst solution.

To immunostain the microglia (Iba1), astrocytes (GFAP), and astrocyte endfeet (AQP-4), the mouse brains were perfused with PBS and 4% PFA, post-fixed in 4% PFA, and dehydrated in 30% sucrose solution. The dehydrated brains were sliced 30 μm thick on a freezing cryostat. Coronal sections were washed 3 times in PBS to completely remove the cryoprotectant solution. After blocking (20% normal donkey serum with 0.1% Triton X-100 in PBS) for 1 hour at room temperature, sections were incubated with primary antibodies (diluted to a ratio of 1:500) for two nights at 4 °C. Secondary antibodies were then added, followed by incubation with Hoechst solution.

To stain neurons (Nissl), the mouse brains were perfused with PBS and 4% PFA, post-fixed in 4% PFA, and dehydrated in 30% sucrose solution. The dehydrated brains were sliced 30 μm thick on a freezing cryostat. Coronal sections were washed in PBS 3 times to completely remove the cryoprotectant solution. Sections were rehydrated in 0.1 M PBS for at least 40 minutes at room temperature. Tissue permeabilization was performed by incubating the sections in PBS containing 0.1% Triton X-100 for 10 minutes, followed by two 5 minutes washes in PBS. NeuroTrace stain was diluted in PBS according to the manufacturer’s recommendations (1:300), and approximately 200 µL of the diluted staining solution was applied to each section to ensure complete coverage. Sections were incubated with the stain for 20 minutes at room temperature. Following staining, sections were washed in PBS containing 0.1% Triton X-100 for 10 minutes, then washed twice in PBS for 5 minutes each. An additional wash in PBS was performed for 2 h at room temperature. Sections were counterstained with Hoechst, rinsed in PBS, and mounted with antifade mounting medium before coverslip placement for imaging.

### *In vivo* toxicity measurement of PLLA-Alb NPs

To investigate the effect of PLLA-Alb NPs, we intravenously injected 150 mg/kg of PLLA-Alb NPs and weighed the mice to record any changes in body weight. Moreover, we performed hematoxylin and eosin (H&E) staining to examine histopathological changes in the major organs of NP-injected and saline-injected mice. After 13 weeks, the mice were transcardially perfused with PBS and 10% Neutral Buffered Formalin (NBF), and the organs were collected and post-fixed in 10% NBF for 24 hours. LSU Diagnostics (Louisiana Animal Disease Diagnostic Lab assisted in H&E staining.

### Fluorescent microscopy, electron microscopy, and imaging analysis

All the fluorescent images were captured using the Olympus SD-OSR spinning disk super-resolution microscope equipped with cellSens Dimension software. The images were acquired using a 4x, 10x, or 100x objective. The TEM images were taken using a JEOL 1400 Biological and Soft Material TEM. The image acquisition settings remained constant throughout the experiment. The image analysis was performed using Fiji/Image-J.

### Statistical Analysis

Statistical analyses were performed using GraphPad Prism 10. All experiments were repeated at least three times. Unpaired *t*-test or One-way ANOVA was performed for statistical analysis. Comparisons among multiple groups were performed using one-way ANOVA followed by Tukey’s multiple- comparison test. Data are presented as means ± SD; where applicable, \**p* < 0.05, \*\**p* < 0.01, \*\*\**p* < 0.001, or \*\*\*\**p* < 0.0001 was considered a statistically significant difference. The specific *N* value per group is provided in the figure captions.

## Acknowledgments

We thank Dr. Yongchan Kwon, Claire Lanclos, and Kassidy Porche for their help with the Western Blot. Confocal microscopy was performed at the Advanced Microscopy and Analytical Core (AMAC) at Louisiana State University. Illustration figures were created with Biorender.com.

## Author contributions

Authors’ contributions are attributed on the basis of the CRediT author statement. SR: methodology, investigation, validation, formal analysis, data curation, visualization, writing - original draft; US: methodology, investigation; DA: investigation, validation; CA: methodology, investigation, validation, writing - review & editing; AS: investigation; CS: methodology, supervision, resources, project administration, writing - review & editing; EC: methodology, supervision, investigation, visualization, formal analysis, data curation, project administration, funding acquisition, Writing - review & editing, QC: Conceptualization, methodology, investigation, visualization, validation, formal analysis, data curation, supervision, resources, project administration, funding acquisition, writing - original draft, Writing - review & editing.

## Data and materials availability

All data needed to evaluate the conclusions in the paper are present in the paper and/or the

Supplementary Materials.

