## Supplemental Information for "On-demand, reversible blood-brain barrier opening via electrical activation of piezoelectric nanoparticles for targeted brain drug delivery"

*Corresponding:

Elisa Castagnola,

Qi Cai,

**This PDF file includes:**

Figs. S1 to S9; Table S1 to S2

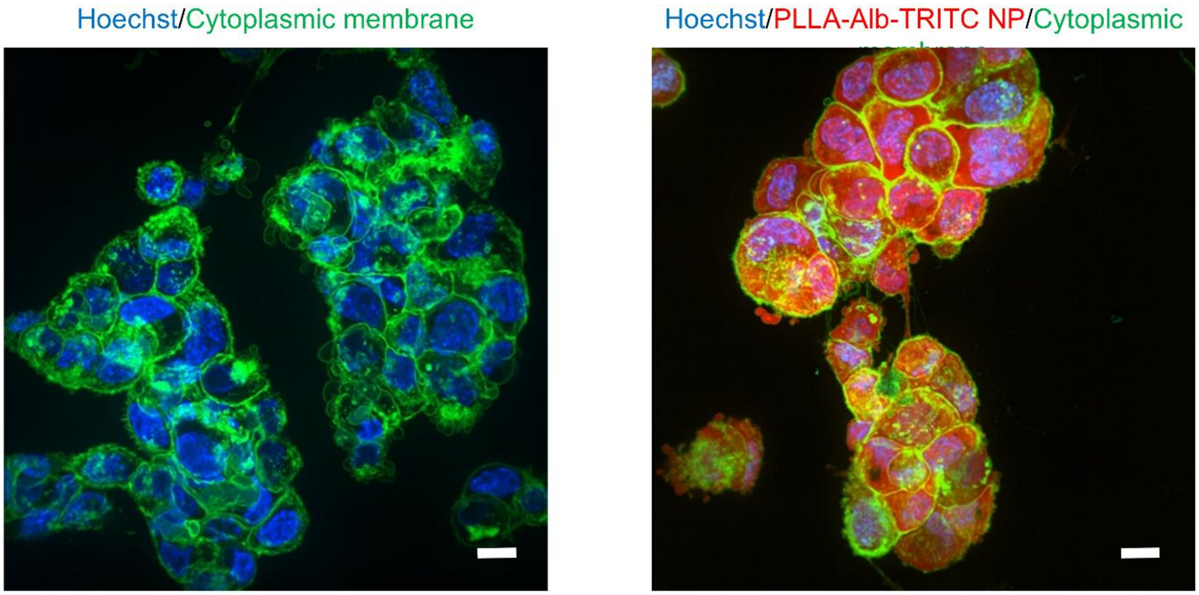


**Fig. S1. Localization of PLLA-Alb-TRITC in human cerebral microvascular endothelial cells.** Scale bar = 10 µm.

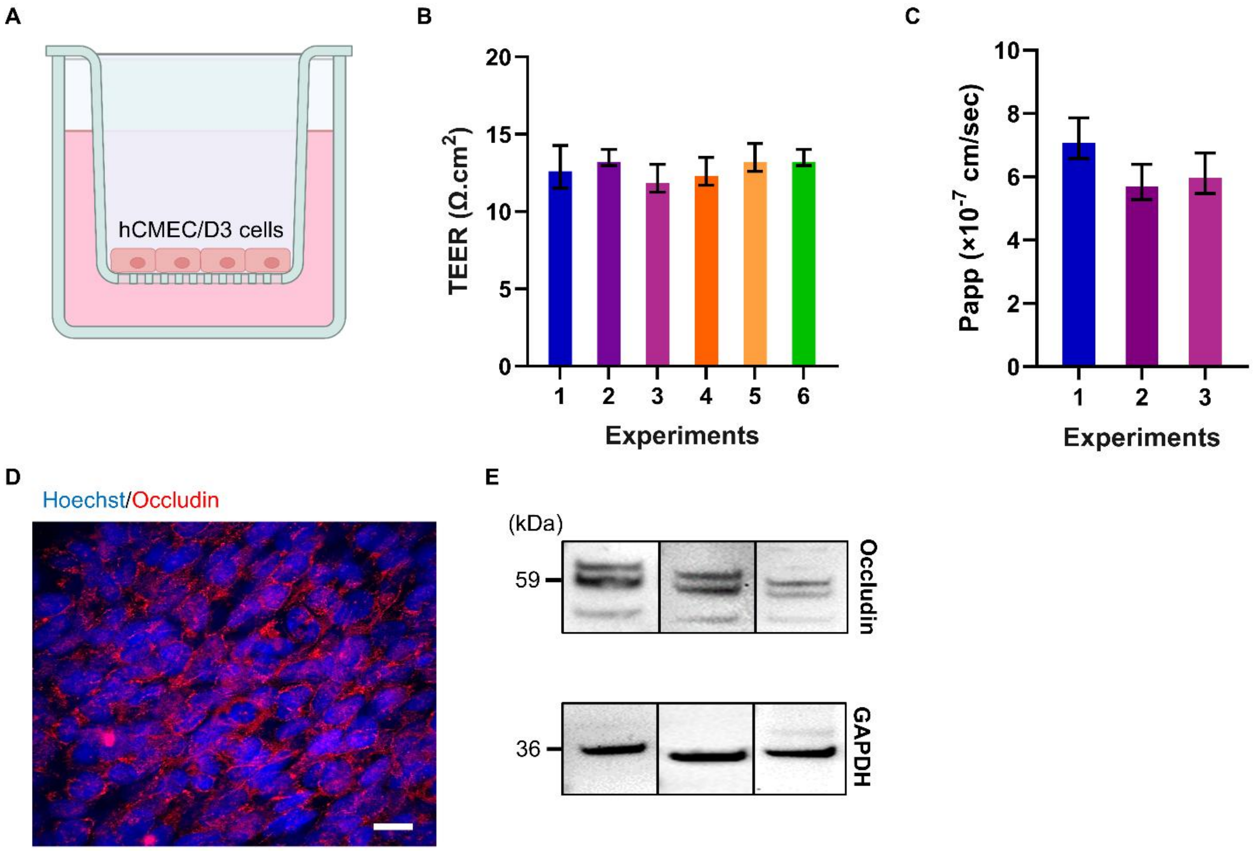


**Fig. S2. Characterization of the in vitro BBB model**. (A) Schematic illustration of the in vitro BBB model. (B) Transendothelial electrical resistance (TEER) measurements of the model. N=3 biological replicates in each experiment. Data are expressed as Mean ± SD. (C) Permeability to FITC-dextran (40 kDa). N=4 biological replicates in each experiment. Data are expressed as Mean ± SD. (D) Occludin formation assessed by immunocytochemistry. Scale bar = 10 µm. (E) Occludin expression confirmed by Western blot.

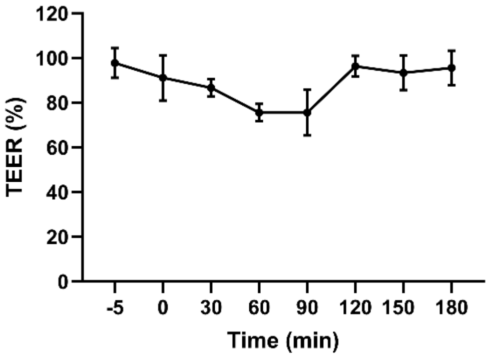


**Fig. S3. Incubation of human cerebral microvascular endothelial cells with BAPTA-AM (10 µM) for 1 hour does not significantly decrease TEER in the *in vitro* BBB model**. N=3 biological replicates. Data are expressed as Mean ± SD. One-way ANOVA, no significant difference.

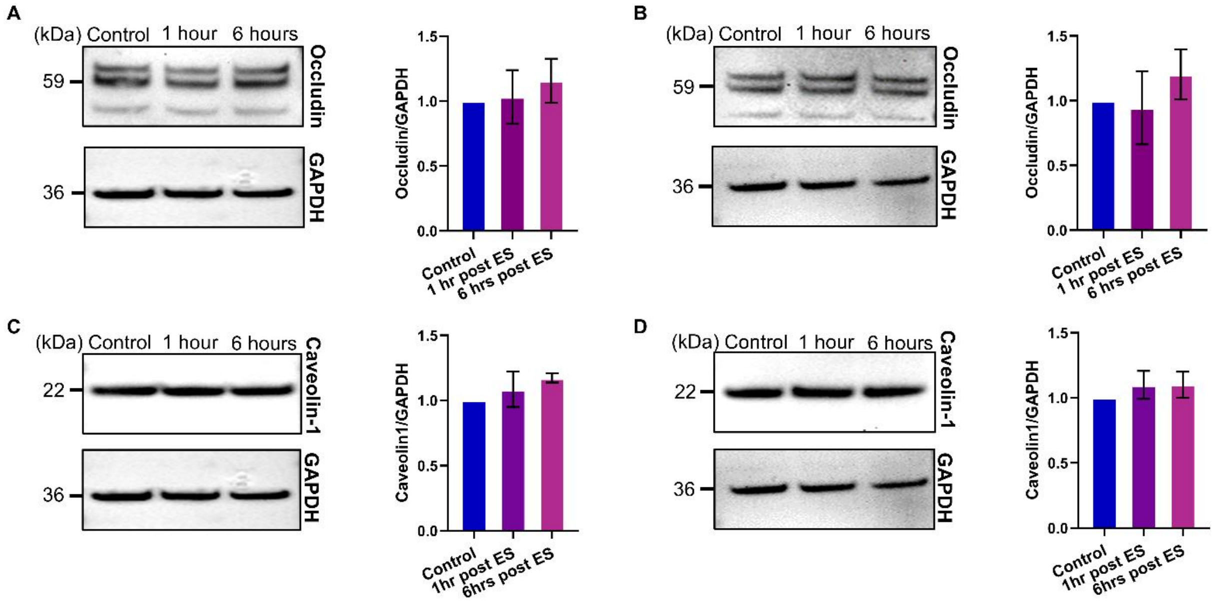


**Fig. S4. Evaluation of the expression of occludin and caveolin-1 under control conditions**. (A) Occludin expression before and after electrical stimulation in the in vitro BBB model using hCMEC/D3 cells. (B) Occludin expression before and after eBBB induced with PLLA-PEG NPs. (C) Caveolin-1 expression before and after electrical stimulation in the in vitro BBB model. (D) Caveolin-1 expression before and after eBBB induced with PLLA-PEG NPs. N=3 biological replicates. One-way ANOVA, no significant difference.

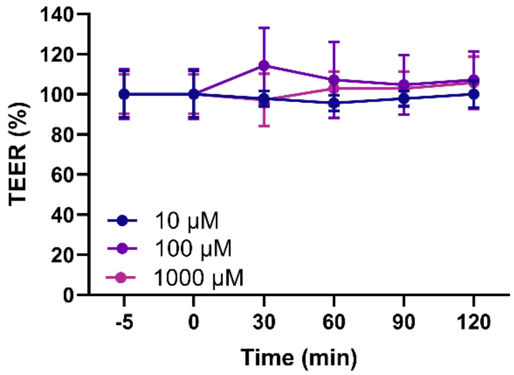


**Fig. S5. Incubation of human cerebral microvascular endothelial cells with mβCD (10-1000 μM) for 1 hour does not significantly decrease TEER in the in vitro BBB model**. N=3 biological replicates. Data are expressed as Mean ± SD. One-way ANOVA, no significant difference.

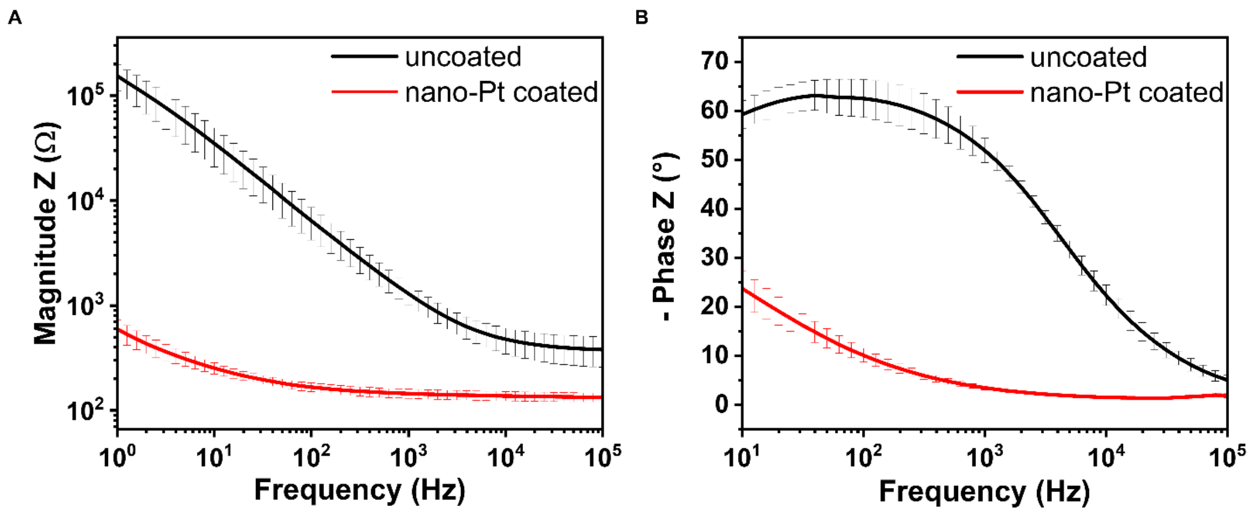


**Fig. S6. Electrochemical impedance spectroscopy (EIS) of 4×1 microelectrode arrays (MEAs) before and after nano-Pt coating, demonstrating reduced electrode impedance following nano-Pt deposition.** (A) Impedance magnitude and (B) phase angle spectra.

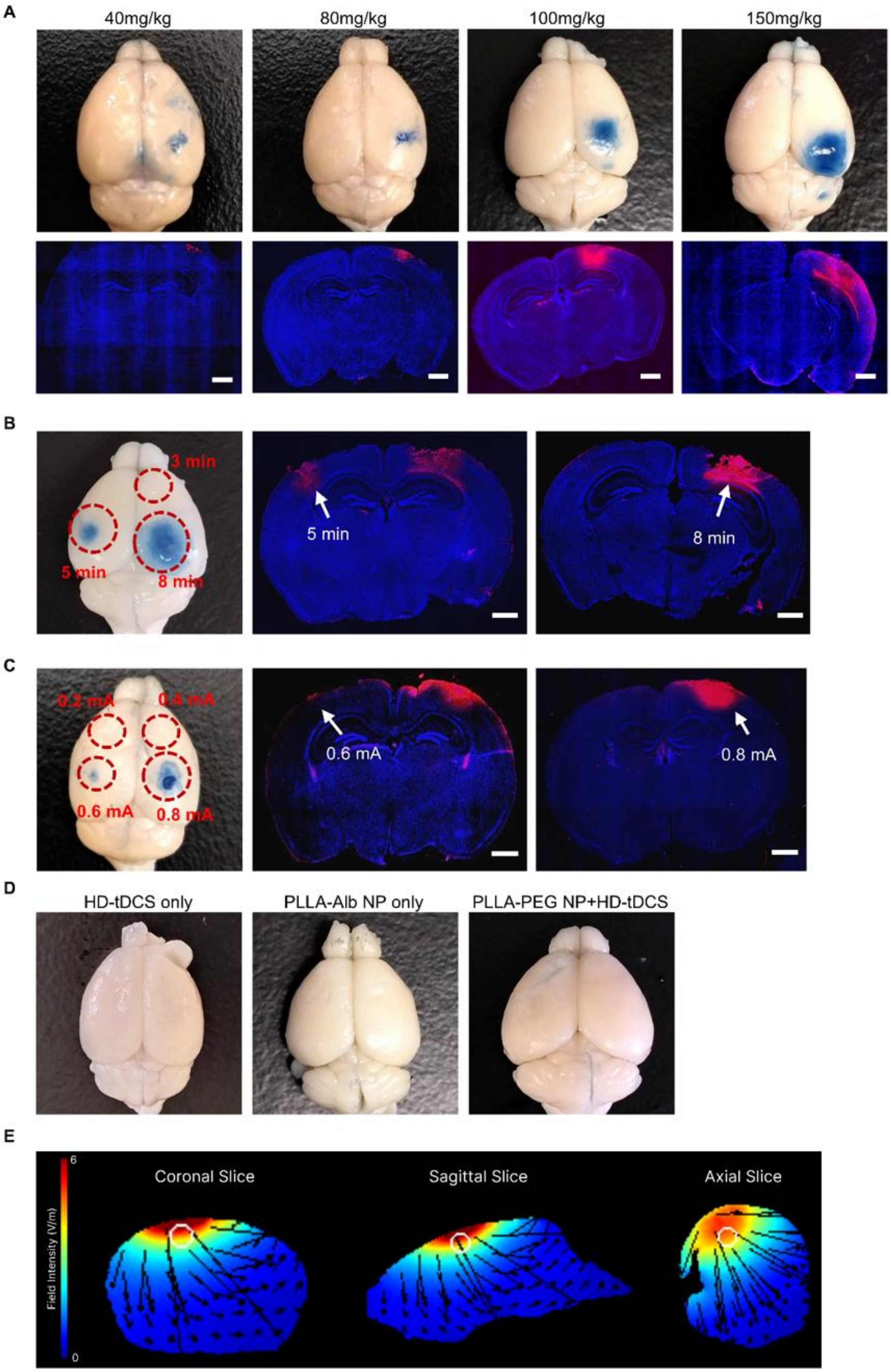
A
40mg/kg
80mg/kg
100mg/kg
150mg/kg
B
3 min
5 min
8 min
5 min
8 min
C
0.2 mA
0.4 mA
0.6 mA
0.8 mA
0.6 mA
0.8 mA
D
HD-tDCS only
PLLA-Alb NP only
PLLA-PEG NP+HD-tDCS
E
Field Intensity (V/m)
6
0
Coronal Slice
Sagittal Slice
Axial Slice

**Fig. S7. eBBB modulation in the mouse brain**. (A) Optimization of the nanoparticle dose. Scale bar= 2 mm. (B) Optimization of the stimulation time. Scale bar = 2 mm. (C) Optimization of the current. Scale bar = 2 mm. (D) Electrical stimulation alone, PLLA-Alb NP injection alone, or electrical stimulation of PLLA-PEG NP does not induce BBB opening. (E) Simulation of electrical field distribution in the mouse brain.

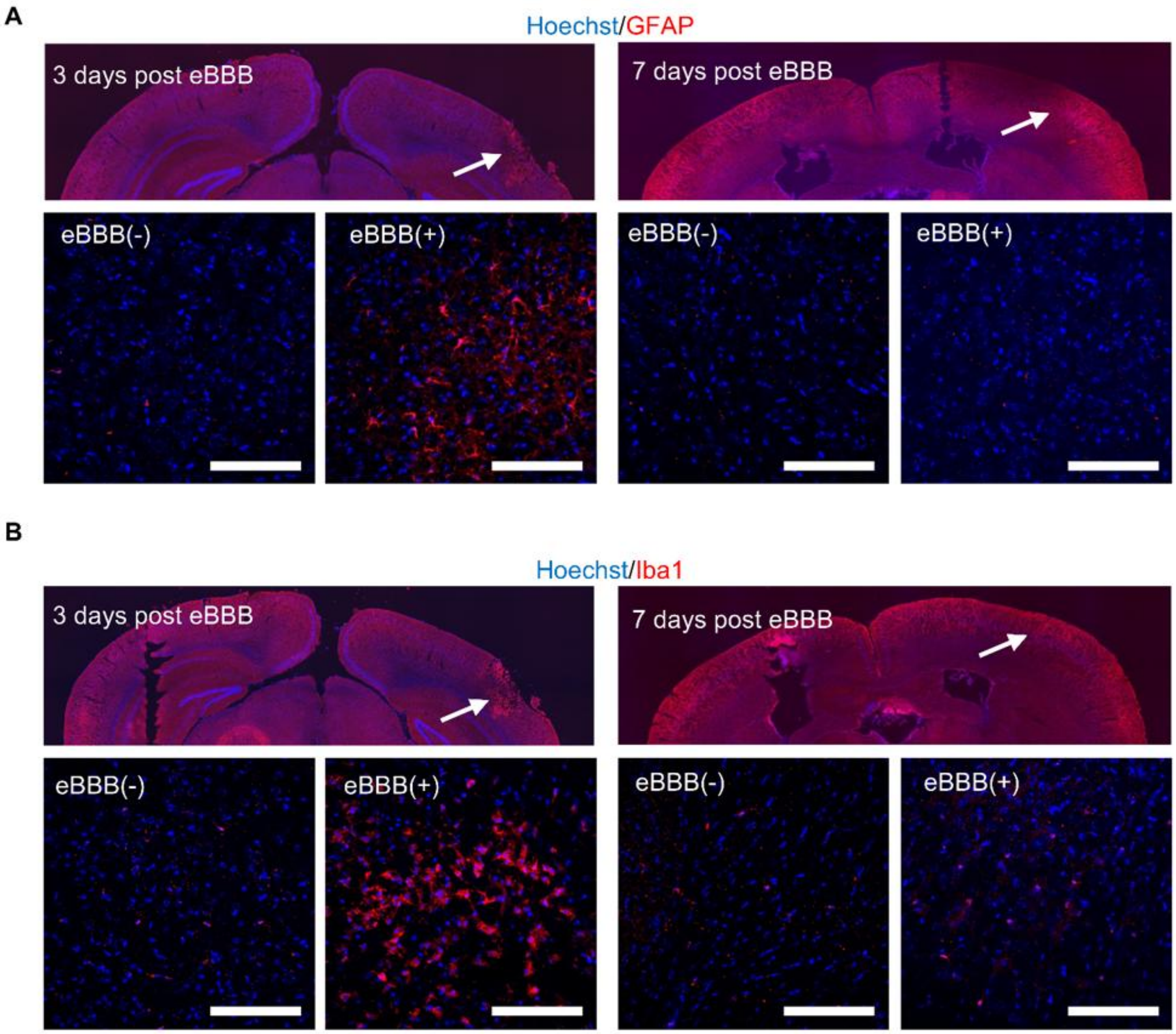


**Fig. S8. eBBB induces a transient glial response.** (A) IHC staining of GFAP$^+$ astrocytes at 3 and 7 days after eBBB modulation shows a transient increase at 3 days that returns to baseline by 7 days. (B) IHC staining of Iba1$^+$ microglia at 3 and 7 days post eBBB reveals a similar transient increase that resolves by 7 days. Scale bar = 100 μm. Arrows indicate the BBB modulation region.

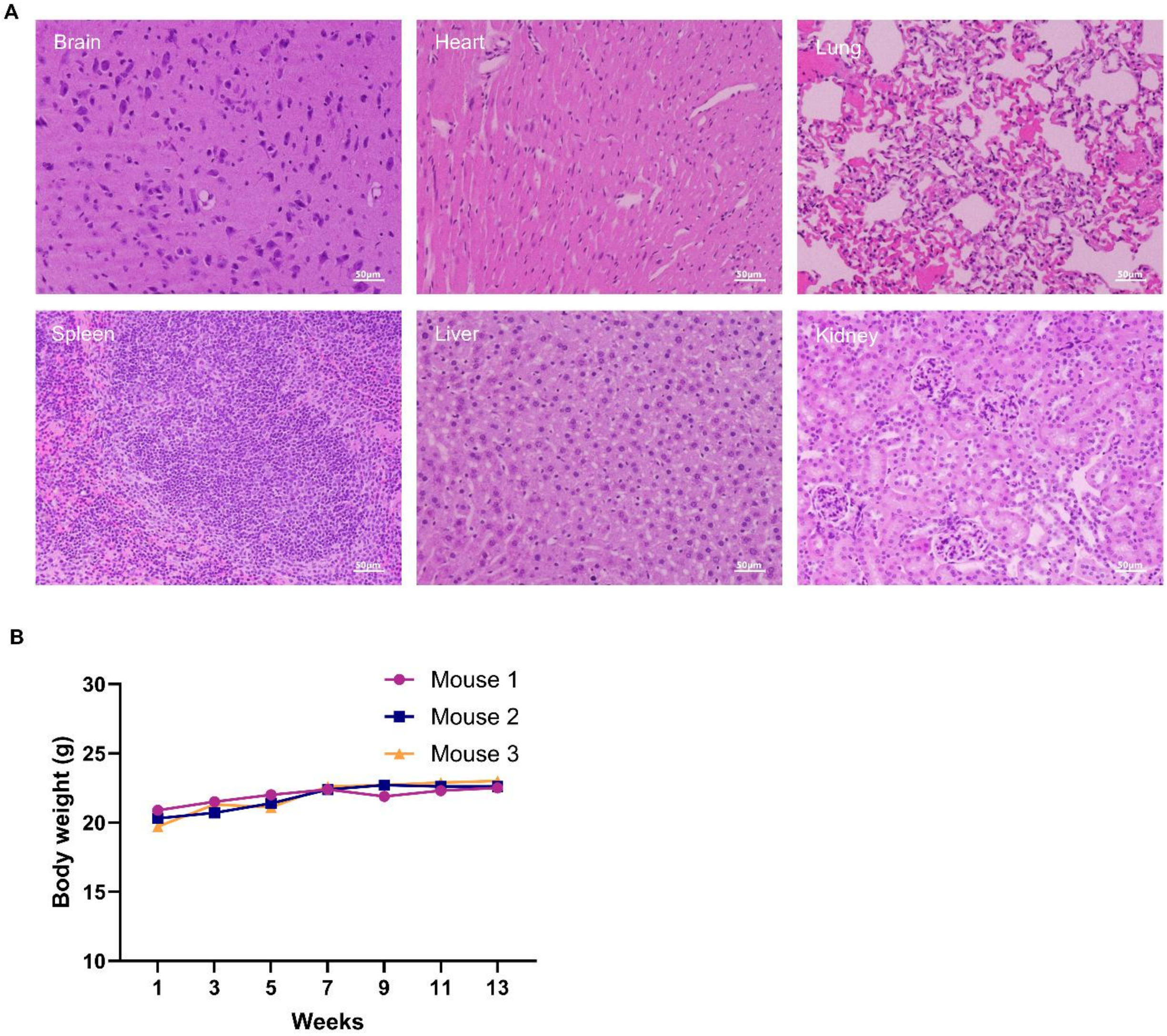


**Fig. S9. Long-term systemic safety of PLLA-Alb nanoparticles**. (A) Hematoxylin and eosin (H&E) staining of major organs harvested 13 weeks after intravenous administration of PLLA-Alb NPs (150 mg/kg). No histological abnormalities, inflammatory infiltrates, or tissue necrosis were observed in any organ. Scale bar = 50 μm. (B) Body weight of mice monitored biweekly over 13 weeks following nanoparticle injection, showing no significant decrease, confirming the absence of systemic toxicity.

**Table S1. PLLA conjugate synthesis conditions.**

| Conjugate | PLLA solution | Oxalyl chloride | DMF | Coupling component |
|---|---|---|---|---|
| **PLLA–Alb** | 2 g in 50 mL DCM | 43 µL | 5 mL | Alb, 2 g in 50 mL anhydrous DMSO |
| **PLLA–PEG** | 1 g in 30 mL DCM | 15 µL | 5 mL | $NH_2$–PEG–CM, 1 g in DCM |
| **PLLA-TRITC** | 1 g in 30 mL DCM | 15 µL | 5 mL | TRITC, 22 mg in anhydrous DMSO |

**Table S2. PLLA nanoparticle formation conditions**

| Formulation | Polymer feed | Organic phase | Aqueous phase | Additive |
|---|---|---|---|---|
| **PLLA–Alb NP** | 350 mg PLLA–Alb | 16 mL DCM | 75 mL water | Trehalose, 2:1 w/w |
| **PLLA–PEG NP** | 450 mg PLLA–PEG | 6 mL DCM | 70 mL water | PVA, 45 mg |
| **PLLA–Alb–TRITC NP** | 350 mg PLLA–Alb + 35 mg PLLA–TRITC | 16 mL DCM | 75mL Water | Trehalose, 2:1 w/w |
| **PLLA–PEG–TRITC NP** | 450 mg PLLA–PEG + 50 mg PLLA–TRITC | 6 mL DCM + 4 mL ethyl acetate | 70 mL water + 10 mL ethyl acetate | PVA, 45 mg |
